# L1 insertion intermediates recombine with one another or with DNA breaks to form genome rearrangements

**DOI:** 10.1101/2025.09.17.676864

**Authors:** Carlos Mendez-Dorantes, Jupiter C. Kalinowski, Cheuk-Ting Law, Phillip Schofield, Aidan Burn, Kathleen H. Burns

## Abstract

LINE-1 retrotransposition is common in human cancers and rearrangements at insertion sites can contribute to cancer-driving oncogene amplifications and promote genome instability. However, the mechanisms underlying rearrangements of L1 retrotransposition intermediates are poorly understood. To address this gap, we developed GFP-based recombination reporter assays to study the formation of L1 retrotransposition-mediated rearrangements. Using these reporters combined with long-read sequencing approaches, we find that L1 retrotransposition intermediates can recombine with distal DNA breaks to generate chromosomal rearrangements. We also find that two distinct L1 insertion intermediates can recombine with each other to generate chromosomal rearrangements. Both types of rearrangements depend on L1-encoded ORF2p endonuclease and reverse transcriptase activities. Using these reporters, we discover that L1 retrotransposition-mediated rearrangements are robustly induced when the recombining sequences share homology and that their formation requires the homologous recombination factor BRCA1. Given the repetitive nature of our genome, these findings highlight the risk of L1 insertion intermediates becoming substrates for aberrant recombination and promoting genome instability.

## Introduction

A third of our genome is composed of repetitive DNA elements derived from the activity of retrotransposons (Hoyt et al. 2022; Mendez-Dorantes and Burns 2023). Retrotransposons are mobile DNA sequences that copy and paste via RNA intermediates and include LINE-1 (L1), *Alu* and SVA in humans. L1 is the only active retrotransposon that encodes its own protein machinery for retrotransposition: ORF1p, an RNA binding protein (Khazina et al. 2011), and ORF2p, a reverse transcriptase (RT) with endonuclease (EN) activity (Mathias et al. 1991; Feng et al. 1996; Baldwin et al. 2024; Thawani et al. 2024). While 17% of our genome is L1 sequence, only about 150 full-length, retrotransposition-competent L1 loci are present in any individual’s genome, and these are commonly silenced in somatic cells. In contrast, L1 ORF1p overexpression is detected across cancers of many distinct tissues of origin (Rodic et al. 2014; Taylor et al. 2023). In these malignancies, L1 retrotransposition is now well appreciated as a mutagenic signature (Lee et al. 2012; Helman et al. 2014; Rodriguez-Martin et al. 2020).

Major sequencing efforts have mapped somatically-acquired L1 insertions in cancer genomes in search of cancer-driving insertional mutations (Lee et al. 2012; Helman et al. 2014; Tubio et al. 2014; Flasch et al. 2019; Rodriguez-Martin et al. 2020) and revealed that L1 insertional mutations crippling tumor suppressor genes are recurrent but uncommon (Scott et al. 2016; Cajuso et al. 2019) . In addition to these canonical insertions, L1 retrotransposition can cause alterations at the insertion site including large segmental deletions leading to the loss of tumor suppressor genes (Gilbert et al. 2002; Symer et al. 2002; Gilbert et al. 2005; Rodriguez-Martin et al. 2020). The recent application of long-read sequencing to cancer genome analyses and to experimental models of L1 overexpression has expanded on our knowledge of the types of alterations caused by L1 retrotransposition (Mendez-Dorantes et al. 2024; Zumalave et al. 2024), revealing more evidence for chromosomal translocations, inversions and complex rearrangements as outcomes of L1 mutagenesis. However, the precise mechanisms underlying L1 retrotransposition-mediated rearrangements remain poorly understood.

Key insights into mechanisms of L1 retrotransposition have been enabled by the development of a cell-based functional assay for retrotransposition (Moran et al. 1996; Ostertag et al. 2000; Liu et al. 2018), which has been a guiding experimental tool in the L1 biology field for the past twenty-five years. In this assay, a reporter gene cassette (e.g., neo, GFP) interrupted by an intron is introduced in antisense orientation in the 3’ UTR of a L1 sequence. The reporter is only activated upon L1 transcription (leading to intron splicing) and genomic insertion (placing the cassette in the genome for expression). When the L1 sequence is intact, this tool largely reports the occurrence of canonical target-primed reverse transcription (TPRT) reactions. L1 insertions preferentially occur at endonuclease cut sites (3’-AA/TTTT-5’), contain polyA sequences at their 3’ end, and are flanked by short, target site duplications (TSDs). Here, we develop novel reporter assays which depend on L1 retrotransposition-mediated rearrangements to reconstitute split GFPs. We validate these as tools to enumerate recombination between L1 insertion intermediates, both with one other or with distal chromosomal breaks. In both instances, recombinations are promoted by sequence homology and host factors that mediate homologous recombination (HR). Our work highlights the labile nature of L1 retrotransposition intermediates and their risk to genome integrity.

## Results

### L1 insertion intermediates recombine with DNA breaks to generate L1 retrotransposition-mediated chromosomal rearrangements

During L1 retrotransposition, reverse transcription is initiated at the 3’-OH of a nick in the genomic target site DNA (Fig 1A). A working model for the formation of a retrotransposition-mediated rearrangement is that the cDNA intermediate fails to resolve to form a double stranded, integrated L1 insertion, but rather the nicked target is converted to a double-stranded DNA break (Fig 1A). This L1-mediated DNA break is then subject to ligation to a distal independent chromosomal break to cause a rearrangement. To directly test this model, we established a novel GFP-based reporter to signal recombination between a L1 retrotransposition intermediate and a distal DNA double strand break (DSB). To do this, we modified the classic L1-GFP retrotransposition reporter assay (Fig 1A). Briefly, the L1-GFP reporter expresses a L1 sequence (L1RP) containing a GFP-AI reporter in the 3’ UTR (Ostertag et al. 2000; Kopera et al. 2016). The reporter consists of a GFP expression cassette that is in the anti-sense direction relative to the L1 sequence and that is interrupted by a sense-oriented intron and flanked by constitutive promoter and polyA signal (Fig 1A). Accordingly, transcription and intron splicing, followed by retrotransposition of the reconstituted GFP cassette into the genome can mark in cells with acquired insertions based on GFP expression. Using this reporter, we found robust L1 retrotransposition in U2OS cells, which was drastically reduced with mutations in the reverse transcriptase (RT) domain and the endonuclease (EN) domain of ORF2p (Fig 1B-C, SFig1A), as previously shown (Mathias et al. 1991; Feng et al. 1996; Tao et al. 2022).

**Figure 1.**
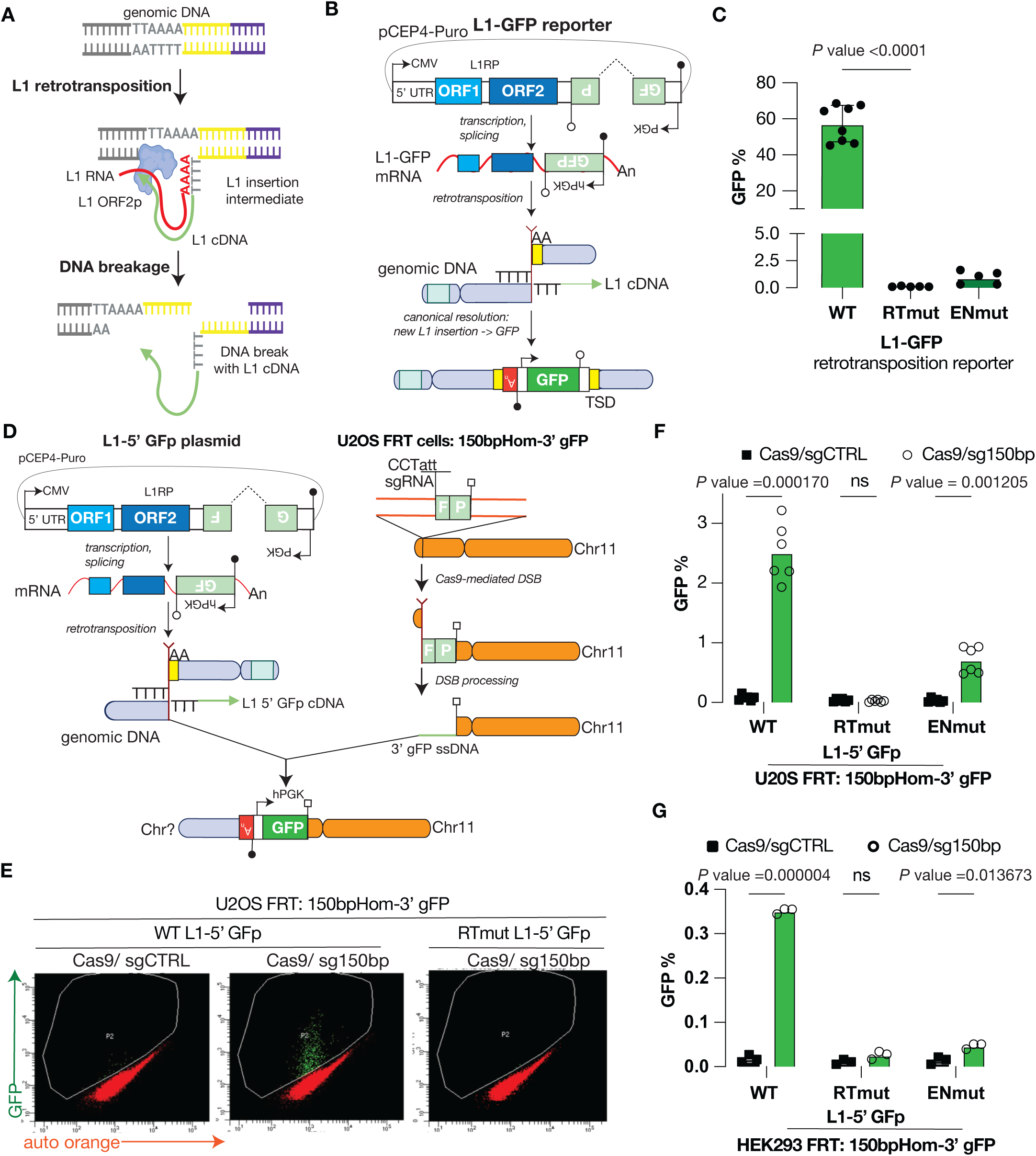
L1 insertion intermediates recombine with DNA breaks to generate L1 retrotransposition-mediated chromosomal rearrangements. **A.** A working model for DNA breakage resulting from L1 retrotranspostion. L1 retrotransposition occurs via target-primed reverse transcription. L1 ORF2p can nick genomic DNA at an EN target sequence (3’-AA/TTTT-5’) to liberate a polyT sequence that complementary bind the L1 RNA polyA sequence to form a primer-template structure. L1 ORF2p can then polymerize from the free 3’OH and use the L1 RNA as a template to generate *de novo* L1 cDNA that can be processed into a double-stranded insertion in the genome. Alternatively, DNA breakage can occur during the resolution of the L1 cDNA intermediate into an insertion generating recombinogenic DNA break ends. **B.** A schematic of the classic L1-GFP reporter assay. A pCEP4-Puro plasmid constitutively expresses a native human L1 sequence (L1RP) expressing ORF1p (RNA binding protein) and ORF2p (endonuclease and reverse transcriptase) containing a GFP reporter in the 3’ UTR that marks *de novo* L1 insertions. The GFP cassette is in the anti-sense direction relative to L1 sequence and is interrupted by an intron in the sense direction. The continuous GFP sequence is generated when the L1 transgene is transcribed and spliced but cannot be expressed and translated until the cassette is retrotransposed. Accordingly, GFP expression marks cells with de novo L1 insertions. **C.** Shown is the retrotransposition frequency of WT, reverse transcriptase mutant (RTmut, D702Y) RTmut or endonuclease mutant (ENmut, H230A) L1 measured by the L1-GFP reporter assay in U2OS cells. n = 6. P-value <0.0001, each compared to RTmut using one-way ANOVA with Dunnett’s test. **D.** Shown is a schematic of a new GFP reporter assay modeling the recombination between a L1-cDNA intermediate and a chromosomal break to generate a chromosomal rearrangement. *Left:* A pCEP4-Puro plasmid contains a modified version of the classic L1-GFP reporter construct, where the 3’ UTR contains an anti-sense 5’ GFP fragment (5’GFp) that is interrupted by a sense intron (L1-5’ GFp plasmid). Transcription, splicing and retrotransposition of the L1 RNA results in a canonical insertion containing an incomplete GFP cassette and does not induce GFP expression. Nonetheless, retrotransposition of the L1 RNA generates L1 insertion intermediates containing an 5’GFP fragment with the potential to be resolved in non-canonical events. *Right:* A 3’ GFP fragment containing 150 bp homology to the 5’ GFP fragment from L1-5’ GFp plasmid (3’ gFP 150bp Hom) was introduced chromosomally in U2OS cells using the FRT/ Flp recombinase system (150bpHom-3’ gFP U2OS cells). A chromosomal break can be induced by the CRISRP/Cas9 system at the edge of the 150bp homology of the 3’gFP 150bp Hom sequence (Cas9/sg150bp). Hence, L1 cDNA insertion intermediates containing a 5’ GFP fragment can recombine with a broken chromosome containing a 3’ GFP fragment forming a chromosomal rearrangement and restoring GFP expression. **E.** Shown are representative flow cytometry plots of 150bpHom-3’ gFP U2OS cells transfected with WT or RTmut L1-5’ GFp reporter plasmid, and Cas9/ sgCTRL or Cas9/ sg150bp expressing plasmid. GFP expression is only induced with transfection of the WT L1-5’ GFp reporter plasmid and the Cas9/150bp expressing plasmid. **F.** Shown is the GFP frequency induced in U2OS FRT 3’ gFP (150bp Hom) cells transfected with WT, RTmut or ENmut L1-5’ GFp reporter plasmid, and Cas9/sgCTRL or Cas9/sg150bp expressing plasmid. n = 6. P-value calculated using unpaired two-tailed Student’s t tests with the Holm-Sidak correction for multiple comparisons. **G.** Shown is the GFP frequency induced in HEK293 150bpHom-3’ gFP cells transfected with WT, RTmut or ENmut L1-5’ GFp reporter plasmid, and Cas9/sgCTRL or Cas9/sg150bp expressing plasmid. n = 3. P-value calculated using unpaired two-tailed Student’s t tests with the Holm-Sidak correction for multiple comparisons.

To establish a GFP reporter for L1 retrotransposition-mediated rearrangements, we first modified the GFP-AI reporter in the 3’ UTR of L1 such that retrotranspositon would generate insertions containing an incomplete GFP sequence (5’ GFP fragment or GFp) (Fig 1D). For this, we introduced an antisense 5’ GFP fragment interrupted by a sense intron into the 3’ UTR of L1 in place of the GFP-AI reporter (L1-5’GFp reporter). In this scenario, retrotransposition of the L1 reporter transcript would be predicted to result in insertions with an incomplete GFP sequence. Importantly, we found that the new L1-5’GFp reporter plasmid expresses ORF1p and ORF2p comparably to the L1-GFP reporter plasmid (SFig 1B). Using the FRT/Flp recombinase system in U2OS cells (Kelso et al. 2019), we generated a cell line containing a chromosomal copy of the remaining 3’ GFP sequence (gFP, Fig 1D, SFig 2). Notably, in this design, we used a 3’ GFP sequence containing 150 bp of overlapping homology with the reconstituted 5’ GFP sequence from the L1-5’GFp reporter: 150bpHom-3’ gFP cells. In addition, we placed a unique target sequence to introduce a DNA double-strand break (DSB) using Cas9/sgRNA (Cas9/sg150bp) at the edge of the 3’ GFP sequence, which we validated by assaying the frequency insertion/deletion (indel) mutations at the targeted site when the Cas9 and the targeting guide were supplied (SFig 3). Using these experimental tools, we found that GFP expression can be induced after co-transfection of the L1-5’GFp reporter plasmid and the Cas9/sg150bp plasmid in 3’ gFP-150bpHom cells (Fig 1E-F), but not with transfection of the L1-5’GFp reporter alone. This suggested that the GFP-based assay could capture recombination events between L1 retrotransposition cDNA intermediates and a broken chromosome and that the DSB was necessary for efficient recombination. Notably, the frequency of GFP was 22.8-fold lower for the GFP recombination reporter versus the canonical L1 GFP retrotransposition reporter (2.5% versus 57%).

**Figure 2.**
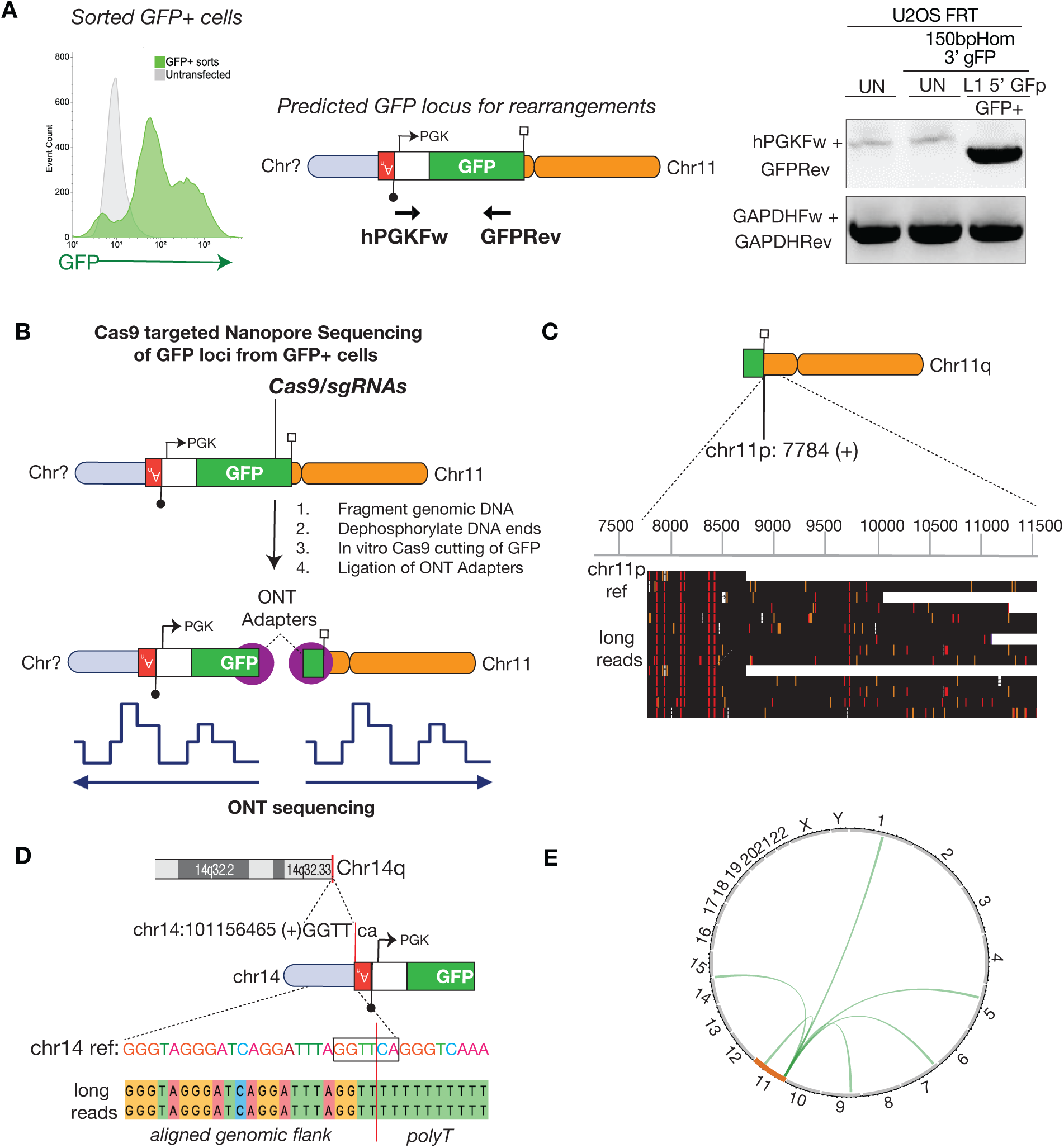
Cas9 targeted nanopore sequencing reveals sequence structure of L1 retrotransposition-mediated rearrangements. **A.** GFP+ cells harbor the predicted recombination product. Shown are PCR amplification products from untransfected cells and sorted GFP+ cells derived from U2OS 150bpHom-3’ gFP cells transfected with WT L1 5’ GFp plasmid and Cas9/sg150bp expressing plasmid using primers hPGKFw (L1 5’GFp plasmid) and GFPrev (3’ gFP 150bp Hom). Amplification of GAPDH locus was used as a control. **B.** Schematic of the Cas9 targeted nanopore sequencing strategy to map the upstream and the downstream sequences of the GFP loci of sorted GFP+ cells. Genomic DNA was isolated from sorted GFP+ cells derived from U2OS FRT 3’ gFP (150bp Hom) cells after transfection with WT L1 5’ GFp plasmid and Cas9/sg150bp expressing plasmid. Genomic DNA was first mechanically fragmented to 30 kb length and dephosphorylated. Dephosphorylated genomic DNA was then incubated with Cas9 ribonucleoproteins using sgRNAs targeted to the 3’gFP sequence at the FRT locus. Cleaved genomic DNA ends are then ligated with Oxford Nanopore Technologies (ONT) sequencing adapters. These DNA fragments were sequenced on a MinION flow cell. Long reads were first mapped to a GFP reference to select GFP-containing long reads for subsequent mapping to the human genome (T2T) to determine their genomic integration site. **C.** Shown are several ONT long reads containing the 3’ GFP fragment that were mapped to the human genome (T2T) revealing the chromosomal FRT locus in U2OS cells is in chromosome 11 (7784 +). **D.** Shown are ONT long reads containing the 5’ GFP fragment and the recombination junction mapping to chromosome 14 (101156465 +) and showing evidence of L1 retrotransposition at the genomic site of integration: inserted at (5’-GGTT/CA-3’) and containing an polyA sequence. **E.** A circos plot showing the genomic locations of L1 insertion intermediates that recombined with the Cas9-mediated DNA double-strand break generated at edge of the 150 bp homology of the 3’gFP (150 bp Hom) locus in chromosome 11 to restore GFP expression via a chromosomal translocation. Each translocation is represented by an arch originating from the p arm of chromosome 11 and terminating at the location of each L1 insertion intermediate.

**Figure 3.**
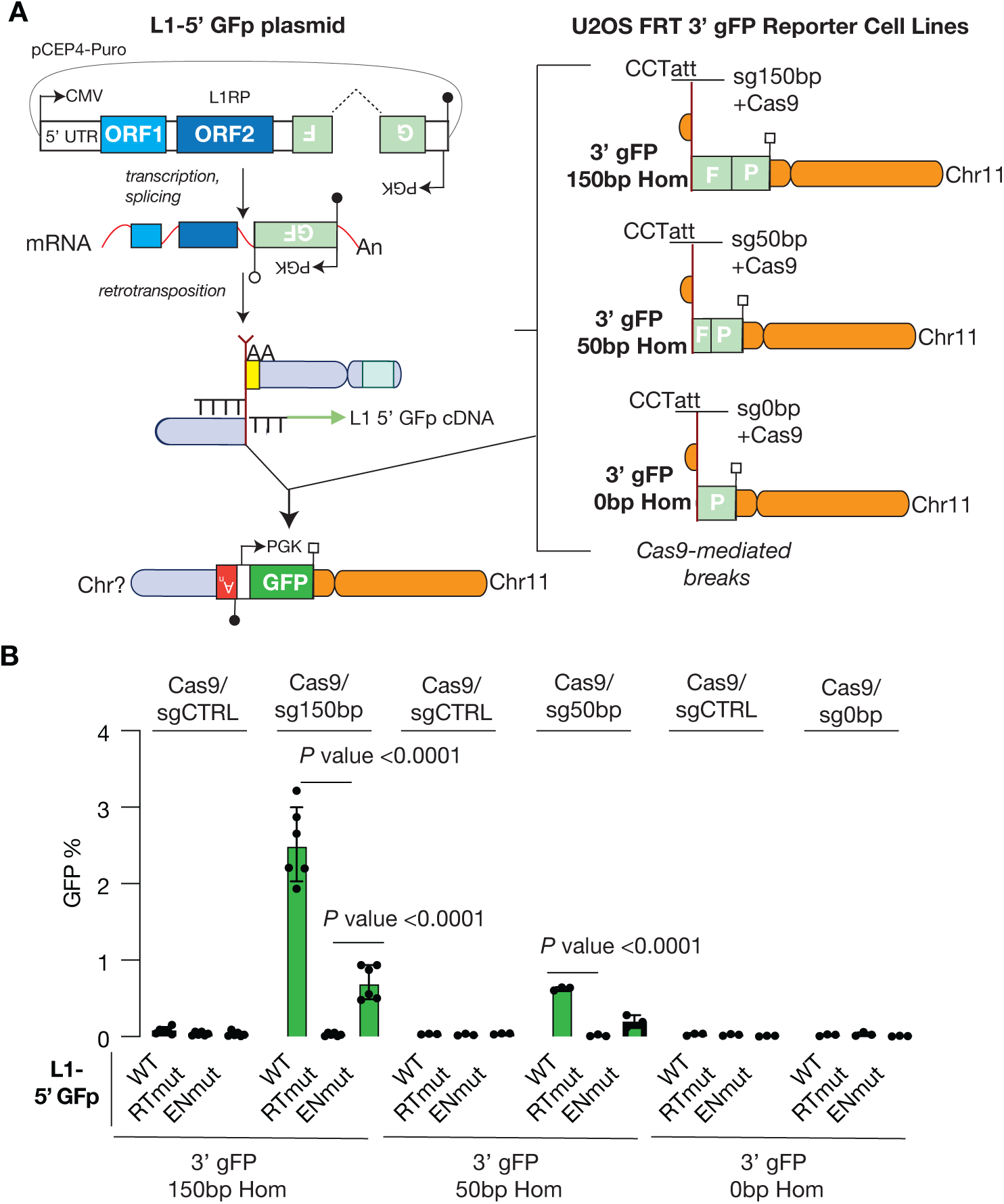
Efficient recombination between L1 reverse transcribed cDNA and chromosomal breaks requires homology to generate chromosomal translocations. **A**. Shown is a schematic of GFP recombination reporters assaying the influence of the length of homologous sequences between a L1-cDNA intermediate and a chromosomal break for the formation of chromosomal rearrangements. *Left:* The L1-5’ GFp plasmid is transfected into cells to generate L1 insertion intermediates containing a 5’GFP fragment with the potential to be resolved in non-canonical events such as rearrangements. *Right:* 3’ GFP fragments containing varying degree of homology to the 5’ GFP fragment from L1-5’ GFp plasmid were introduced chromosomally in U2OS cells using the FRT/ Flp recombinase system: 3’gFP 150bp Hom, 3’gFP 50bp Hom, and 3’gFP 0bp Hom. Each of these 3’GFP fragments was engineered to contain a specific sgRNA target sequence to induce a Cas9-mediated chromosomal break at the edge of the homology. Hence, we can express the L1-5’GFp plasmid and the respective Cas9/sgRNA plasmid in each 3’gFP reporter cell line to monitor the influence of the homology between L1 cDNA insertion intermediates containing a 5’ GFP fragment and a broken chromosome containing a 3’ GFP fragment on the formation of a chromosomal rearrangement by assaying GFP expression. **B.** Shown is the GFP frequency induced in each U2OS 3’gFP reporter cell line transfected with WT, RTmut or ENmut L1-5’ GFp reporter plasmid, and the respective Cas9/sgRNA expressing plasmid or the control Cas9/sgCTRL expressing plasmid. n = 6 or 3. P-value calculated by comparing WT and ENmut versus RTmut in each condition using a two-way ANOVA with the Dunnett correction for multiple comparisons.

We performed several experiments to validate the GFP recombination assay for L1 retrotransposition-mediated rearrangements. For example, we found that mutating the RT domain of ORF2p in the L1-5’GFp plasmid resulted in no GFP+ cells (Fig 1E-F), supporting that ORF2p RT activity is required to make *de novo* L1 cDNA products capable of recombining to restore the GFP cassette. This result also excludes the possibility that that GFP expression is caused by recombination directly between the L1 plasmid containing the interrupted 5’ GFp sequence and the broken chromosome containing 3’gFP sequence. In addition, we found that mutation of the EN domain of ORF2p reduced the frequency of GFP+ cells (Fig 1F), demonstrating that efficient endonuclease activity during TPRT promotes GFP reconstitution. As noted above, we found that expression of the WT L1-5’GFp plasmid alone without induction of the chromosomal break in 150bpHom-3’gFP cells failed to induce GFP+ cells, suggesting that both active L1 retrotransposition and DSB induction are required to produce a functional GFP cassette. We examined this further by performing sequential transfection of the L1-5’GFp reporter plasmid followed by the Cas9/sg150bp plasmid allowing resolved L1 insertions to accumulate in cells prior to inducing the chromosomal DSB (SFig 4). For this, we first selected cells that were transfected with the L1-5’GFp reporter plasmid to allow L1-5’GFp insertions to accumulate but then expanded these cells without selection to dilute the episomal plasmid (SFig 4B). These cells are GFP negative but should contain fully resolved L1 insertions containing 5’GFP sequences, which we used to introduce the the Cas9/sg150bp plasmid simultaneously with an empty vector control, a second WT L1-5’GFp reporter plasmid or a RT mutant L1-5’GFp reporter plasmid (SFig 4C). We found that the chromosomal breakage alone in cells previously exposed to the L1-5’GFp reporter plasmid failed to induce GFP+ cells (SFig 4C). However, transfection of the WT L1-5’GFp plasmid concurrent with induction of the chromosomal break produced GFP expression. This result is consistent with our hypothesis that both active retrotransposition (i.e., unresolved insertion intermediates) and DSB induction are concurrently required to induce a recombination event between them.

**Figure 4.**
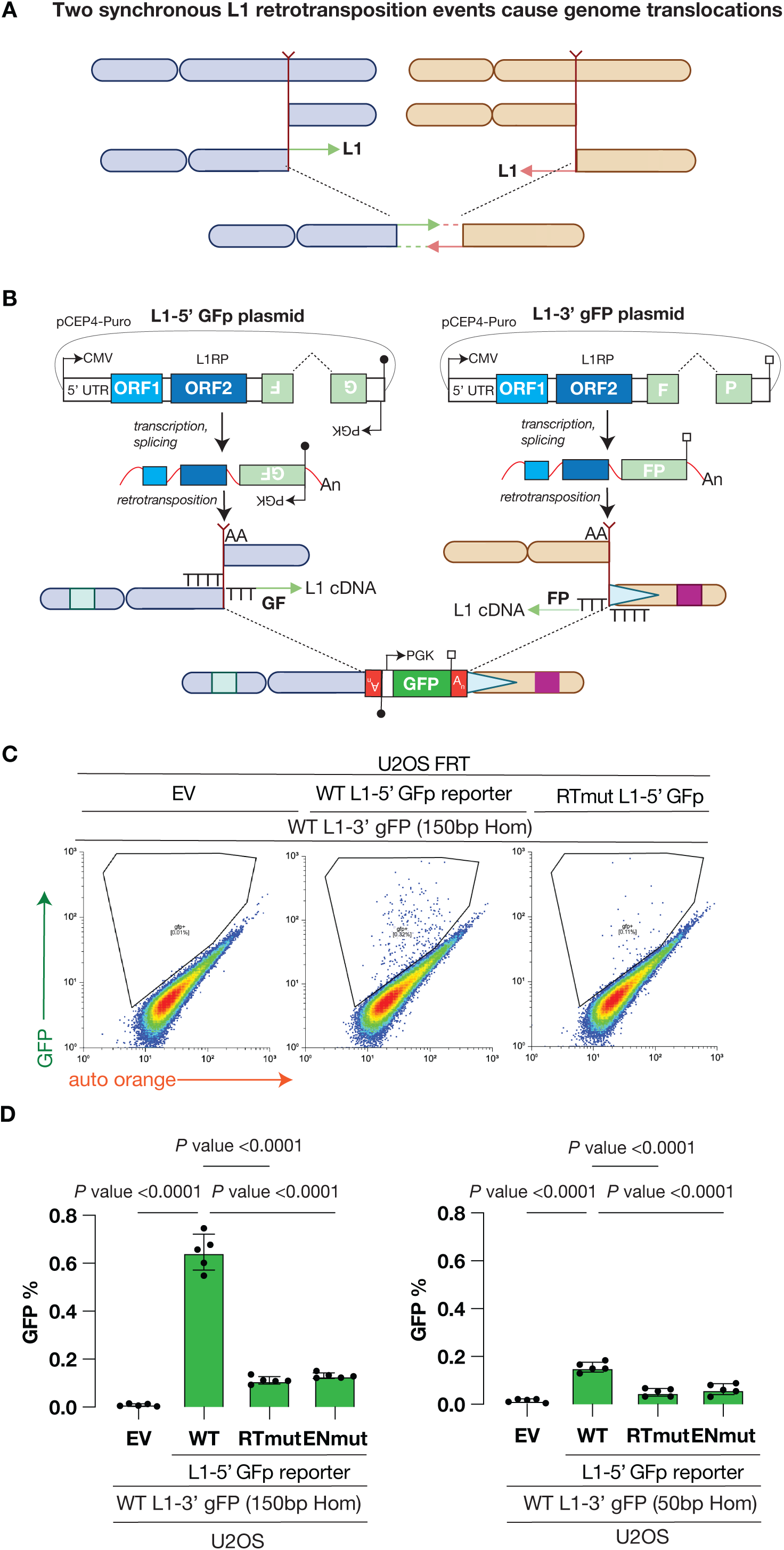
Two L1 retrotransposition events can recombine to cause genome rearrangements. **A.** A working model for how two independent L1 retrotransposition events on distinct chromosomes can cause a chromosomal translocation. **B.** Shown is a novel GFP-based reporter assay to ascertain recombination between two independent L1 retrotransposition intermediates generating a chromosomal rearrangement. *Left:* The L1-5’ GFp plasmid is transfected into cells to generate L1 insertion intermediates containing a 5’GFP fragment with the potential to be resolved in non-canonical events such as rearrangements. *Right:* A pCEP4-Puro plasmid contains a modified version of the classic L1-GFP reporter construct, where the 3’ UTR contains a sense 3’ GFP fragment (3’gFP) that is interrupted by a sense intron (L1-3’ gFP plasmid). Transcription, splicing and retrotransposition of the L1 RNA results in a canonical insertion encompassing an incomplete GFP cassette. Both reporter constructs the L1-5’ GFp plasmid and L1-3’ gFP plasmid can be expressed in cells to generate L1 reverse transcribed cDNA intermediates that share homologous sequences within the GFP fragment which can recombine to generate a GFP-expressing rearrangement. **C.** Shown are representative flow cytometry plots of parental U2OS cells transfected with EV, WT or RTmut L1-5’ GFp reporter plasmid, and the WT L1-3’gFP plasmid, which 3’GFP fragment contains 150bp of homology to the 5’ GFP fragment to L1-5’GFp plasmid. GFP expression is only induced with transfection of the WT L1-5’ GFp reporter plasmid and the WT L1-3’ gFP reporter plasmid. **D.** Shown are the frequencies of GFP+ cells induced in parental U2OS cells transfected with EV, WT, RTmut or ENmut L1 5’GFp plasmid with WT L1-3’gFP (150 bp Hom) or WT L1-3’gFP (50 bp Hom). n=5 P-value calculated using one-way ANOVA test comparing to WT L1 5’ GFp with Hom-Sidak correction for multiple comparisons.

Lastly, we established the GFP-based recombination reporter in HEK293 cells and found similar results (Fig 1G) except the frequency of GFP+ cells was lower as compared to U2OS cells. We found that GFP expression is produced by co-transfection of the L1-5’GFp reporter plasmid and the Cas9/sg150bp plasmid in 150bpHom-3’gFP HEK293 cells (Fig 1G, Fig 2A), and that this is dependent on L1 ORF2p enzymatic activities. These results support the model that retrotransposition intermediates recombine with chromosomal breaks, and that this is not specific to U2OS cells.

### Sequence structures of L1 retrotransposition-mediated genome rearrangements

We next sought to sequence GFP recombination events induced in 150bpHom-3’ gFP U2OS cells transfected with WT L1-5’GFp plasmid and Cas9/sg150bp. For this, we sorted GFP+ cells and assayed for presence of the predicted GFP by PCR; as expected, we found only GFP+ cells, and not control cells, contained the predicted PCR product demonstrating juxtaposition of the 3-phosphoglycerate kinase (PGK) promoter and the 3’ end of the GFP coding sequence (Fig 2A). To sequence these recombination sites and their flanking DNA, we used Cas9 to target Oxford Nanopore Technology (ONT) sequencing to gDNA extracted from the sorted GFP+ cells (Fig 2B). Briefly, we sheared genomic DNA into large fragments, dephosphorylated the ends, and performed *in vitro* Cas9 cutting using sgRNAs targeting the 3’ end of the 3’ GFP sequence inserted at the chromosomal FRT locus of U2OS cells. This approach leaves phosphorylated DNA ends capable of ligation to the nanopore adapter preferentially at the GFP. We ligated adapters and sequenced these by ONT sequencing, kept reads with a GFP sequence, and mapped these to the T2T human reference genome to determine their genomic locations. Using this strategy, we expected that reads aligning to the 3’ end of GFP will map to the chr11 location where the 3’ gFP (150bo Hom) sequence is integrated at the FRT site, whereas reads aligning to the 5’ end of GFP will include the rearrangement junction between the spliced 5’ GFP and 3’ GFP sequences and will extend to distinct chromosomal locations wherever a retrotransposition reaction had been initiated (SFig 5A).

**Figure 5.**
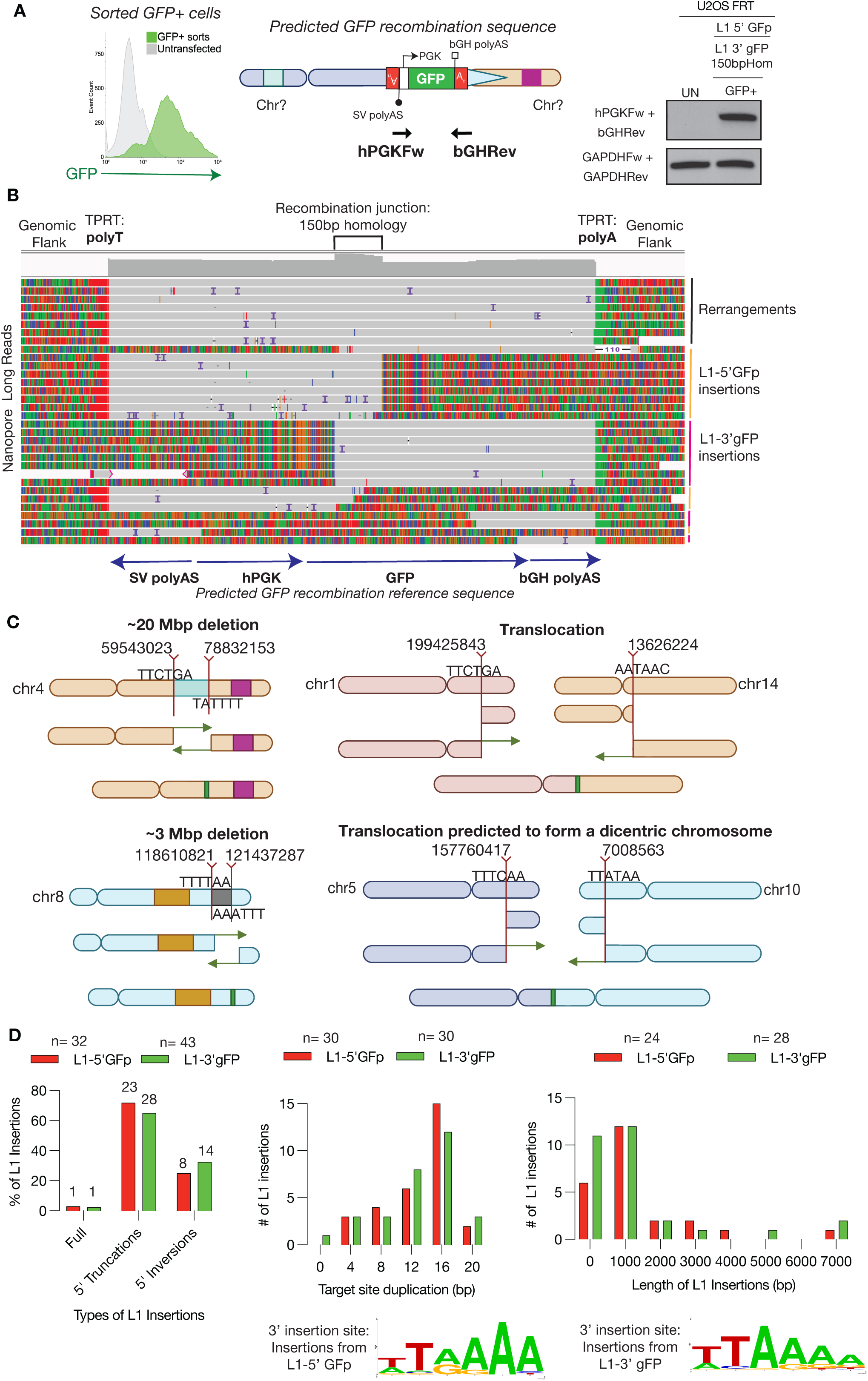
Nanopore whole genome sequencing reveals sequence structure of retrotransposition-mediated rearrangements induced by two independent L1 retrotransposition events. **A.** Sorted GFP+ cells harbor the predicted GFP recombination product. Shown are PCR amplification products from sorted GFP+ cells derived from parental U2OS cells transfected with L1-5’ GFp plasmid and with L1-3’ gFP plasmid, as well as untransfected cells, using primers hPGKFw (L1-5’GFp plasmid) and bGHRev (L1-3’gFP plasmid). Amplification of the GAPDH locus was used as a control. **B.** Representative Nanopore long reads from GFP+ cells that align to the predicted GFP recombination sequence. Three types of aligned reads emerge: Type 1) Rearrangements include reads that span the complete recombination reference sequence and are flanked by L1 TPRT evidence such as inserted polyA sequences at the genomic junctions. Type 2) L1 canonical insertions by L1-5’GFp reporter plasmid include reads that only map to the 5’ side of the recombination reference sequence. Type 3) L1 canonical insertions by L1-3’GFp reporter plasmid include reads that only map to the 3’ side of the recombination reference sequence. **C.** Shown are schematics of the chromosomal rearrangements inferred by reading through the recombined GFP cassette: two segmental deletion rearrangements, a translocation predicted to generate a stable derivative chromosome, and a translocation predicted to form an unstable dicentric chromosome. Shown are the locations of the L1 retrotransposition events in each chromosome and the target sequences that resembled the EN cleavage motif providing evidence of L1 TPRT activity. **D.** Landscape of L1 insertions in GFP+ cells induced by the L1-5’ GFp reporter plasmid and the L1-3’ gFP reporter plasmid. Shown are the number of L1 insertions, including full-length insertions, 5’ truncations and 5’ inversions. Also, shown are the length of target site duplications of L1 insertions and the length of L1 insertions generated by L1-5’GFp and L1-3’gFP. Finally, also shown are the logo sequence motif of the genomic sequence where L1 insertions were integrated at the 3’ end, reconstituting the known EN cleavage motif sequence: 5’-TT/AAAA-3’.

Consistent with this model, we found that long reads aligning to the 3’ end of GFP all mapped to the subtelomeric region of Chr11p arm: chr11:7784 (Fig 2C, Table 1), where the FRT construct was introduced. Moreover, we found reads aligning to the 5’ end of the GFP sequence extended from GFP into sequence mapping to several distinct chromosome locations (Fig 2C-E, SFig 5A-B, Table 1). Importantly, all these reads contained the rearrangement junction between the spliced 5’ GFP sequence and the 3’ GFP sequence, the hPGK promoter, and the SVpolyA signal (Fig 2D, SFig 5A-B, Table 1). As expected, L1 sequence is lost in these recombination events. Notably, the detection of spliced 5’GFP sequence in these reads provides evidence of the spliced L1 RNA as an intermediate for the reconstitution of GFP, consistent with the requirement of ORF2p RT activity (Fig 1F-G). Consistent with this result, the 5’ GFP reads reflect signatures of L1 TPRT including inserted polyA sequences upstream of the SV polyA signal and, in three cases, an EN cleavage motif (5’ nT/AAAn 3’). In total, this strategy allowed us to sequence six chromosomal rearrangements with five interchromosomal translocations and one inversion rearrangement in Chr11 (Fig 2E, Table 1). Lastly, we also captured two events showing evidence of L1-5’GFp inserting at the targeted Cas9-mediated break in Chr11p arm and recombining with the proximal 3’gFP (SFig. 5C, Table 1). We previously reported that L1 can cause insertions at Cas9-mediated DNA breaks (Tao et al. 2022), which is consistent with L1 exploiting pre-existing DNA lesions (Morrish et al. 2007). Together, these sequencing data support that GFP expression induced by our recombination reporter models L1 retrotransposition-mediated chromosomal rearrangements.

### L1 retrotransposition-mediated genome rearrangements are promoted by sequence homology

Homologous sequences are known to mediate the formation of chromosomal rearrangements in cancer genomes (Mendez-Dorantes et al. 2018; Li et al. 2020). To address the influence of length of homology between L1-reverse transcribed cDNAs and DSB ends on the frequency of their recombination, we engineered two additional 3’ gFP cell lines. Each was designed to harbor a chromosomal copy of the 3’ gFP sequence with less overlapping homology to the 5’ GFp cDNA, 50 bp or 0 bp (producing 50bpHom-3’ gFP cells and 0bpHom-3’ gFP cells) (Fig 3A, SFig 2B, SFig 3). We then transfected WT L1-5’GFp plasmid into each of the series of 3’gf/FP cell lines while inducing chromosomal breaks at the integrated 3’ reporter sites and compared the frequencies of resulting GFP+ cells. We found that reducing the overlapping homology from 150 bp to 50 bp reduced the frequency of GFP recombination events by 4-fold (Fig 3B), demonstrating that homologous sequences between L1 cDNA intermediates and chromosomal breaks promote their recombination. In support of this, we found that cells with 0bp of homology failed to convert to GFP+ after expression of the L1-5’GFp plasmid and the induction of the chromosomal break in the 0bpHom-3’gFP cell line (Fig 3B). As an important control, we tested the efficiency of each sgRNA to induce Cas9 cutting at the target sequence of each reporter cell line by quantifying the frequency of insertion/deletion mutations and found that each sgRNA induced a comparable frequency of mutagenesis (SFig 3). In addition, we examined the influence of inducing a Cas9-mediated chromosomal break using the same sgRNA targeting an I-*Sce*I site that we introduced located 12 bp away from the 3’GFP sequences in all reporter cell lines (SFig 6A). We found that induction of a Cas9-mediated chromosomal break using the same sgRNA across all reporter cell lines resulted in a similar trend of GFP+ cells with 50 bp versus 150bp of homology reducing the frequency of GFP+ cells and 0 bp failing to induce GFP+ cells (SFig 6B). This revealed that inducing a chromosomal break 12 bp away from the homology versus at the edge of the homology reduced the frequency of GFP+ cells despite efficient I-*Sce*I DSB induction assayed by indel analysis (SFig 6C,SFig 6D), which is consistent with previous reports showing that the proximity of DSBs relative to the homologous recombining sequences promotes their recombination (Mendez-Dorantes et al. 2018; Kelso et al. 2019; Mendez-Dorantes et al. 2020). Finally, we found that GFP+ cells induced by 50 bp of shared homology between the L1 cDNA intermediates and the chromosomal break were also dependent on the L1 ORF2p enzymatic activities (Fig 3B, SFig 6). Together, these results show that the length of homology between L1 cDNA intermediates and chromosomal breaks impacts the frequency of genome rearrangements.

**Figure 6.**
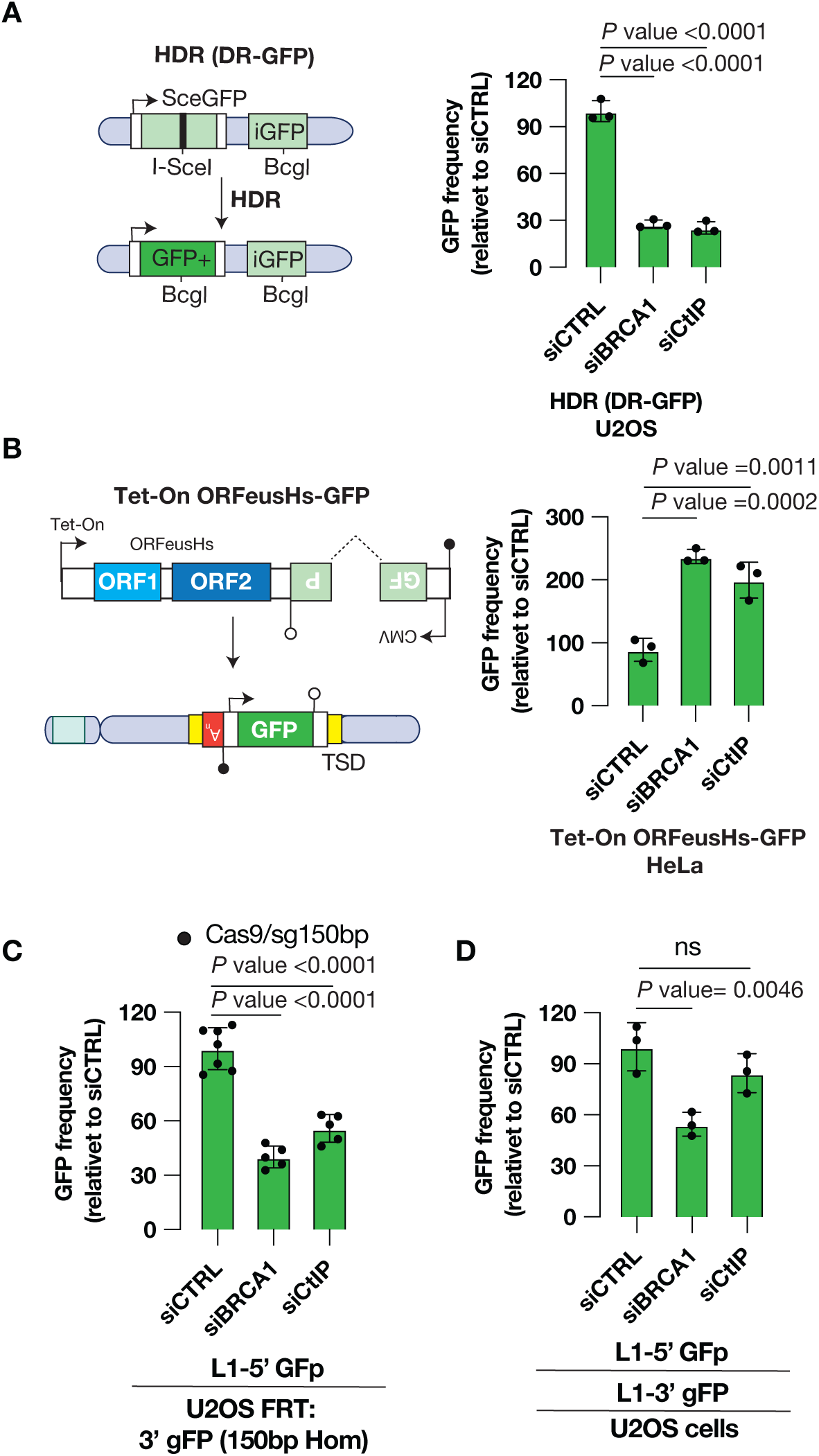
BRCA1 inhibits L1 retrotransposition but mediates L1 retrotransposition-mediated rearrangements dependent on homologous sequences. **A.** BRCA1 and CtIP mediate homologous recombination. Shown is a schematic of the homology-directed repair (HDR) DR-GFP reporter assay chromosomally integrated in U2OS cells. Also, shown are the frequency of GFP+ cells after induction of the DR-GFP reporter assay in cells treated with siCTRL, siBRCA1 or siCtIP. n=3. P-value calculated using one-way ANOVA with Dunnett’s test for multiple corrections. **B.** BRCA1 and CtIP suppress L1 retrotransposition. Shown is a schematic of the Tet-On ORFeusHs-GFP reporter integrated on HeLa cells, which upon doxycycline treatment the L1-GFP reporter transcript is expressed to generate *de novo* L1 insertions marked by GFP expression. Shown is the frequency of L1 retrotransposition in cells treated with siCTRL, siBRCA1 or siCtIP. n=3. P-value calculated using one-way ANOVA with Dunnett’s test for multiple corrections. **C.** BRCA1 and CtIP mediates recombination between L1 cDNA intermediates and chromosomal breaks with homologous sequences for genome rearrangements. Shown are the frequencies of GFP+ cells in 150bpHom-3’gFP U2OS cells treated with siCTRL, siBRCA or siCtIP and co-transfected with WT L1-5’GFp reporter plasmid and the Cas9/sg150bp plasmid. n=7 for siCTRL and n=5 for siBRCA1 and siCtIP. P-value calculated using one-way ANOVA with Dunnett’s test for multiple corrections. **D.** BRCA1, but not CtIP, appears to promote recombination between L1 cDNA intermediates that share homologous sequences. Parental U2OS cells were treated with siCTRL, siBRCA, or siCtIP and co-transfected with the L1-5’GFp plasmid and the L1-3’gFP reporter plasmid. n=3. P-value calculated using one-way ANOVA with Dunnett’s test for multiple corrections.

### Two retrotransposition intermediates can recombine to induce genome rearrangements

Using L1 overexpression systems in combination with whole genome sequencing, we recently reported that two independent, synchronously occurring retrotransposition events can recombine to produce genome rearrangements (Mendez-Dorantes et al. 2024), a phenomenon also recently observed in cancer genomes with high rate of L1 retrotransposition (Zumalave et al. 2024). A working model is that two L1 cDNA products generated by two independent TPRT reactions can recombine to bridge two distant chromosome breaks and thus mediate a long-range genome rearrangement (Fig 4A). To test this model and develop a strategy to enumerate these events, we designed an additional construct for use in combination with our L1-5’GFp reporter that would generate insertion intermediates containing a 3’ gFP sequence (L1-3’gFP reporter plasmid, Fig 4B). Used together, these two split reporters would have the potential to produce cDNAs that in turn may recombine to generate a complete GFP expression cassette (Fig 4B). Given the directionality required, we designed the L1-3’ gFP reporter to contain a sense oriented 3’ GFP fragment interrupted by an intron in the same orientation. Alone, the incomplete GFP fragment cannot be translated into functional protein from the spliced L1 transcript; nor is it functional from a retrotransposed copy. However, we hypothesized that simultaneous TPRT of both the L1-5’GFp transcript and the L1-3’gFP transcript would generate reverse transcribed cDNA intermediates with the potential to recombine to reconstitute the GFP cassette via genome rearrangements (Fig 4B). We made two variants of the L1-3’gFP reporter plasmid, one containing 150bp or another containing 50bp of overlapping homology with the L1 cDNA products generated by the L1-5’GFp reporter plasmid. Using these reagents, we found that co-transfection of the L1-5’GFp plasmid and the 150bpHom-L1-3’gFP plasmid in parental U2OS cells induced GFP+ cells (Fig 4C, Fig 4D), supporting our model that recombination between two *de novo* cDNA products can recombine to restore GFP expression. We found that reducing the overlapping homology from 150bp to 50bp between the two L1 cDNA products resulted in a reduction of GFP+ cells (Fig 4D), demonstrating that homologous sequences between two L1cDNA products promotes their recombination. We also found that mutations in the RT or EN domain of ORF2p expressed from the L1-5’GFp reporter plasmid significantly reduced the frequency of GFP+ cells (Fig. 4D), demonstrating that efficient cDNA production via TPRT is required to restore GFP expression. This result also excludes the possibility that GFP expression in this system is driven by recombination between the two plasmids. Interestingly, we found that the WT ORF2p encoded by the L1-3’gFP plasmid mostly failed to *trans* complement the RT mutant ORF2p encoded by the L1-5’GFp plasmid, which is consistent with the known *cis* preference of ORF2p to associate with its own RNA (Wei et al. 2001; Kulpa and Moran 2006). As a control, we found that both L1-5’ GFp plasmid and L1-3’ gFP plasmid express comparable levels of ORF1p and ORF2p (SFig 7A). Together, these results support our model that two L1 reverse transcribed products can recombine to restore GFP expression.

We then sought to corroborate this interpretation by determining the sequences of GFP recombination events generated by the dual L1 reporter assay. For this, we transfected both the L1-5’GFp reporter plasmid and the L1-3’gFP reporter plasmid with 150bp of homology into U2OS parental cells and we then sorted to enrich for GFP+ cells (Fig 5A). We first assayed for the predicted GFP recombination junction by PCR analysis and found that only GFP+ cells, but not control cells, contained the predicted single PCR product (Fig 5A). To map the recombination events in the genome, we performed ONT whole-genome sequencing of the sorted GFP+ cells, as previously described (Jain et al. 2018). We identified GFP-containing nanopore long reads by aligning reads to the predicted GFP recombination reference sequence and aligned the selected reads to the T2T human genome reference to determine their genomic locations (Fig 5B). We found long reads that spanned the complete GFP sequence (perfect reconstitution events) contained evidence supporting chromosomal rearrangements mediated by recombination between cDNA products of TPRT (Fig 5B, Table 2). For one, we found that the upstream and downstream sequences of the reconstituted GFP sequence mapped to two distinct chromosomal locations (Fig 5B, Table 2), supporting our model that the reconstitution of GFP was caused by genome rearrangements. In addition, we found clear genomic evidence of L1 TPRT activity at the 5’ and 3’ genomic junctions including inserted polyA sequences at sequences that resembled the EN cleavage motif (Fig 5B, Fig 5C, Table 2). Moreover, the evidence of the spliced 5’GFP sequence and the spliced 3’GFP sequence without introns in these reads also provides evidence of the spliced L1 RNAs being involved in the reconstitution of GFP during TPRT, consistent with a requirement of ORF2p RT activity (Fig 4D). Together, these results support our model that two independent products of TPRT at distinct chromosomes recombine to produce GFP expressing genome rearrangements.

In total, we detected five chromosomal rearrangements reconstituting a GFP expression cassette at the breakpoint junction (Fig 5C, Table 2). We found two large segmental deletions on chr 4 and chr8, and a translocation between chromosome 1 and chromosome 14 resulting in a predicted stable der(1;14) chromosome. We also found evidence of a translocation between chromosome 5 and chromosome 10 which would be predicted to result in an unstable dicentric chromosome (Fig 5C, Table 2). In addition, we found two examples of L1 insertions that reconstituted the GFP recombination sequence occurring on the same chromosomal integration site (SFig 7B, Table 2). These events were likely generated by two RT reactions: 1) L1-3’gFP initiated TPRT to generate the 3’GFP cDNA product and resulted in DNA breakage and 2) L1-5’GFp generated 5’GFP cDNA product from the free DNA end break and two cDNA products recombined to seal the insertion in the genome (SFig 7B, Table 2). This model is supported by the fact that both two events contained target site duplications and inserted polyA sequences (Table 2). Together, these results shown that our dual L1 GFP reporter assay models L1 retrotransposition-mediated rearrangements.

Lastly, our approach to apply nanopore WGS to cells transfected with L1 reporter plasmids allowed us to capture retrotransposition products generated by the L1-5’ GFp reporter and L1-3’ gFP reporter plasmids. We found that reads that partially aligned to GFP reflected canonical L1 insertions generated by each respective reporter: reads that only mapped to the 5’ side of the GFP sequence largely represent canonical insertions generated by the L1-5’GFp reporter plasmid while reads that only mapped to the 3’ side of the GFP represented canonical insertions generated by the L1-3’gFP reporter plasmid (Fig 5B, Table 3). In total, we found 75 canonical L1 insertions with 32 insertions made by the L1-5’GFp reporter plasmid and 43 insertions made by the L1-3’gFP reporter plasmid (Fig 5D, Table 3). We found similar frequency of full-length insertions (3.1% versus 2.3), 5’ truncations (71.9% versus 65.1%) and 5’ inversions (25% versus 32.6%) between the two L1 expression constructs. Importantly, these L1 insertions contained variable short target site duplications (TSDs) and were found enriched in the EN cleavage motif sequence: 5’-TT/AAAA-3’ (Fig 5D, Table 3), consistent with genomic signature of L1 TPRT activity. Consistent with this, we found all insertions contained spliced GFP sequences providing evidence that the L1 RNA transcripts were reversed transcribed into cDNAs and inserted into the genome via L1 ORF2p. Finally, we found that most insertions were 5’ truncated resulting in insertions shorter than 2kb as previously reported (Mendez-Dorantes et al. 2024). Together, these results confirm that our L1 reporter plasmids that we modified to develop GFP-based recombination reporters recapitulate known features of *de novo* L1 insertions.

### *BRCA1* mediates L1 retrotransposition-mediated rearrangements dependent on homologous sequences

We and others previously showed that cellular factors involved in homologous recombination such as BRCA1 and the end resection factor CtIP suppress L1 retrotransposition (Liu et al. 2018; Ardeljan et al. 2020; Mita et al. 2020). We hypothesized that BRCA1 and CtIP may similarly suppress L1 retrotransposition-mediated rearrangements. On the other hand, the fact that homology promotes these rearrangements might predict a role for BRCA1 and CtIP. To investigate this, we used siRNAs targeting BRCA1 (siBRCA1) and CtIP (siCtIP) to deplete each respective factor from cells, which we validated by confirming that siBRCA1 and siCtIP treatment reduced the efficiency of homologous recombination measured using the homology-directed repair reporter DR-GFP (Fig 6A) (Moynahan et al. 1999; Stark et al. 2004). Consistent with previous reports, we also found that depletion of BRCA1 or CtIP resulted in robust increase in L1 retrotransposition using a classic retrotransposition reporter (Fig 6B, SFig 8), demonstrating that BRCA1 and CtIP are indeed suppressors of L1 retrotransposition.

In contrast to the suppressive role of BRCA1 and CtiIP on canonical L1 retrotransposition, we find that depletion of BRCA1 or CtIP resulted in a reduction of GFP recombination events induced between L1 cDNA products and chromosomal breaks that share 150bp of overlapping homology (Fig 6C). An explanation for a role for BRCA1 and CtIP in these GFP recombination events is that the chromosomal DSB end needs to be processed into single stranded DNA to reveal homologous ssDNA that can then be recombined with L1 cDNA substrate. Concordant with a role for BRCA1 in such a recombination, we found that depletion of BRCA1 results in a reduction of GFP recombination events induced by two L1 cDNA products that share 150bp of overlapping homology (Fig 6D). In contrast, we found depletion of CtIP had no significant suppressive effect on the frequency of recombination events between two L1 cDNA products that share homology. It is possible that CtIP is dispensable in this scenario since end resection is not required for the recombining of two reverse transcribed single stranded cDNA substrates. Together, these results demonstrate that BRCA1 promotes L1 retrotransposition-mediated rearrangements dependent on homologous sequences.

## Discussion

L1 retrotransposition is common in human cancers and L1 activity can be a source of chromosomal rearrangements and genome instability as demonstrated in experimental studies and cancer genome analyses (Gilbert et al. 2002; Symer et al. 2002; Rodriguez-Martin et al. 2020; Mendez-Dorantes et al. 2024; Zumalave et al. 2024). Here we develop and validate novel GFP recombination reporter assays to study non-canonical resolutions of L1 retrotransposition intermediates into chromosomal rearrangements. In doing so, we provide experimental evidence supporting two models L1 retrotransposition-mediated chromosomal rearrangements. In the first, we show that TPRT cDNA products can recombine with distal DSBs to generate genome rearrangements. In the second, we see that L1 retrotransposition intermediates generated at two independent locations can recombine with each other to juxtapose distant sequences. Our approach to enrich for these events using split GFP reporters and our application of long read sequencing in GFP+ cells to corroborate the rearrangements allowed to make these observations, which have been difficult to study given their rarity and limitations of short-read sequencing. Together, our observations support a model wherein L1 retrotransposition intermediates are labile DNA lesions that can be tethered together and incite chromosomal rearrangements, which is consistent with several reports showing L1 expression is genotoxic in cells (Gasior et al. 2006; Ardeljan et al. 2020; Mendez-Dorantes et al. 2024).

One limitation is that these reporters do not reveal simple end joining events but rather depend on reconstitution of an intact GFP reporter cassette. They are uniquely well suited to study roles of homologous recombination in the resolution of L1 insertion intermediates. Mechanistically, we see that longer intervals of homology shared between L1 cDNA products and a DSB elsewhere in the genome can promote recombination between these sequences to generate large-scale rearrangements. This is also true of homology between synchronously occurring L1 cDNAs, which promotes their recombination. Consistent with this, reporter systems for both types of rearrangements (cDNA-DSB and cDNA-cDNA) reveal a role for the homologous recombination factor BRCA1 in the formation of genome rearrangements. We find a reduction of L1 retrotransposition-mediated rearrangements in cells depleted for BRCA1 despite there being more canonical L1 insertions completed in this genetic background. It remains to be address how BRCA1 may mediate these types of rearrangements via aberrant homologous recombination repair including non-conservative repair pathways such as break-induced replication and single-strand annealing (Bhargava et al. 2016; Kramara et al. 2018; Mendez-Dorantes et al. 2018).

There are many mechanistic questions that remain unaddressed, and we have only a partial understanding of roles for L1 ORF2p and host factors in mediating the completion of a canonical retrotransposition event versus a retrotransposition-mediated rearrangement. ORF2p RT has been shown to polymerizes on several substrates *in vitro* including extending a DNA polymerization on a DNA template in addition to reverse transcribing cDNA on an RNA substrate predicted by the TPRT model (Baldwin et al. 2024); whether ORF2p RT extends annealed DNA homologous sequences generated during rearrangement formation remains to be addressed. Here, we report that BRCA1 has opposing roles for L1 retrotransposition versus retrotransposition-mediated rearrangements that use homologous sequences; delineating whether each depends on end processing or other activities is an area for investigation. Lastly, our finding that two concurrently unresolved L1 intermediates can recombine underscores the open question of the number of L1 retrotransposition intermediates occurring in a single cell cycle, the need to detect and define the structures of these labile lesions, and the need to evaluate whether these intermediates are sequestered in three-dimensional space, as well as to understand how these factors affects risk of aberrant repair via genome rearrangements.

Our study has implications in the fields of cancer biology and genome editing. For cancer biology, our work adds to emerging evidence implicating L1 retrotransposition as a significant endogenous contributor to genome instability. Here, we found 5 retrotransposition-mediated rearrangement and 75 canonical L1 insertions, which is close to the ratio of L1 insertions and L1-mediated rearrangements detected in cancer genomes (Zumalave et al. 2024). Given that cancer genomes can harbor hundreds of somatically acquired insertions, this ratio suggests L1-associated rearrangements may be a potent source of instability. For the field of genome editing, many developing CRISPR technologies are using engineered retrotransposons (Manoj et al. 2021; Wilkinson et al. 2023; Wang et al. 2025). Our study highlights potential unintended consequences of having long cDNA products generated by reverse transcriptase such as ORF2p. These could include segmental deletion rearrangements and long-range genome rearrangements that can generate unstable chromosomes. In addition, we previously showed that L1 insertions can occur at Cas9-mediated DNA breaks (Tao et al. 2022), which is an observation also made in our experimental work here; this may represent an alternative to end joining outcomes depending on the level of retrotransposition.

Our strategy to enrich for L1 retrotransposition-mediated rearrangement using GFP recombination assays is powerful for enriching and studying these infrequent but deleterious rearrangements. In the future, we will want to devise strategies to detect L1 retrotransposition-mediated rearrangements that lack homologous sequences, which appear to be more common based on our prior work sequencing many of these breakpoint junctions (Mendez-Dorantes et al. 2024). In those systems, we see L1 cDNA products can be directly ligated to one another or to chromosomal break likely by end joining repair to form genome rearrangements. These are not modeled by GFP recombination reporters described here.

In summary, our observations highlight that L1 activity can be a significant source of genome instability via the recombination of intermediates of retrotransposition. This study describes the first cellular reporters of this phenomenon, which indicate roles for homology in promoting these rearrangements based on their design. We anticipate that long read sequencing of cancer genomes with high rates of retrotransposition will uncover examples supporting homology mediated recombination at insertion sites.

## Methods

### Cell lines

The two parental cell lines for this study are human osteosarcoma U2OS and human embryonic kidney HEK293 cells, both stably containing the Flp-In T-REx established with the pFRT/lacZeo plasmid (Morales et al. 2015; Kelso et al. 2019). The DR-GFP U2OS reporter cell line was a generous gift from Dr. Jeremy Stark (Gunn and Stark 2012). HeLa cells used to make the Tet-On L1-GFP reporter cell line were acquired from ATTC. All cell lines were cultured at 37°C and 5% CO_2_ using DMEM (Gibco) supplemented 10% FBS, 100 IU/ml penicillin, and 100 μg/ml streptomycin (Invitrogen). The Tet-On L1-GFP HeLa cell line was cultured in Tet-Free FBS media.

### Plasmid construction and cell line generation

All plasmids from this study will be deposited in Addgene.

#### pCEP4 L1 expressing plasmids

All L1 reporter plasmids contain the indicated sequences of L1RP: a retrotransposition-competent human L1, GenBank: AF148856.1 (Kimberland et al. 1999). All L1 constructs are under the expression of the CMV promoter in a pCEP4 episomal vector (Invitrogen), which has been modified to contain the resistance gene for Puromycin instead of Hygromycin. All L1 plasmids contain the 5’UTR of L1, ORF1, the inter-ORF spacer, ORF2, the 3’ UTR modified to contain different version of a GFP reporter cassette and the SV40 polyA tail.

#### The L1-GFP reporter plasmid

(pCMD66B) contains a GFP-based retrotransposition reporter in the 3’ UTR, which is an anti-sense GFP cassette that is interrupted by a sense human gamma globin intron (GFP-AI reporter (Kopera et al. 2016)). The GFP-AI reporter contains a human PGK promoter upstream and a HSV TK polyA signal downstream. The human PGK promoter upstream of the GFP-AI reporter was PCR amplified and introduced via Gibson assembly into the previously validated pCEP4-Puro L1RP GFP-AI reporter plasmid containing the GFP cassette under a CMV promoter (pCMD60ZA (Tao et al. 2022)). Retrotransposition-incompetent mutants of the L1-GFP reporter plasmid were generated containing mutants in ORF2, including a reverse transcriptase mutant (pCMD67A, D702Y) and an endonuclease mutant (pCMD68A, H230A), which were also derived from previously validated mutant L1RP GFP-AI reporter plasmids (Tao et al. 2022).

##### L1-5’GFp reporter plasmid

To generate the L1-5’GFp reporter plasmid (pCMD66A) and the L1-3’gFP reporter plasmids (pCMD172A and pCMD173AA), GFP fragments were synthesized via Genewiz or Twist Bioscience to replace the canonical GFP-AI reporter cassette in the 3’ UTR of the L1-GFP reporter plasmid (pCMD66B). Namely, a 5’ GFP fragment interrupted by <u>an anti-</u> <u>sense</u> human gamma globin intron was synthesized and introduced into the L1 3’UTR of L1-GFP (pCMD68A) replacing the canonical GFP-AI cassette to generate the L1-5’GFp reporter plasmid. Transcription and splicing of the L1 RNA from the L1-5’GFp reporter plasmid could generates a L1 RNA transcript with an antisense 5’GFP fragment in the 3’ UTR but its retrotransposition results in a L1 insertion with an incomplete GFP sequence. The 5’ GFP fragment is under the expression of the human PGK promoter. Retrotransposition-incompetent mutants of the L1-5’GFp reporter plasmid were generated containing mutants in ORF2, including a reverse transcriptase mutant (pCMD70C, D702Y) and an endonuclease mutant (pCMD71B, H230A).

##### L1-3’ gFP reporter plasmid

For the L1-3’ gFP reporter plasmids, two 3’ GFP sequence interrupted by <u>a sense</u> human gamma globin intron followed by a bGH polyA signal was synthesized and introduced into the 3’ UTR of the L1-GFP reporter plasmid (pCMD68A) replacing the GFP-AI cassette, the human PGK promoter and the SV polyA signal. Each of these 3’ GFP sequences contain varying degrees of homology to the 5’ GFP sequence from the L1-5’GFp reporter plasmid: L1-3’gFP (150bp Hom) [pCMD173AA] or L1-3’gFP (50bp Hom) [pCMD172A]. Transcription and splicing of the L1 RNA from the L1-3’gFP reporter plasmid could generate a L1 RNA transcript with a sense 3’ GFP fragment in the 3’ UTR but its retrotransposition results in a L1 insertion with an incomplete GFP sequence.

#### pDNA5/FRT plasmid

To establish cell lines containing 3’GFP sequences integrated chromosomally, we used Flp-In T-REx in U2OS and HEK293 cells each containing an FRT site chromosomally integrated. We first generated targeting vectors containing 3’ GFP sequences in front of a BGH polyA signal. For this, 3’ GFP fragments were synthesized via Genewiz and cloned into the pDNA5/FRT targeting plasmid (generous gifts from Dr. Jeremy Stark). The 3’ GFP fragments were designed to contain varying degrees of homology to the 5’GFP fragment in the L1-5’GFp reporter plasmid, including 150 bp (pCMD15C), 50 bp (pCMD14B) and 0 bp. (pCMD12A). These plasmids were then integrated into U2OS or HEK293 Flp-In T-Rex cells by co-transfection with the PGK-Flp recombinase vector (generous gifts from Dr. Jeremy Stark) using FugeneHD (Promega), as previously described (Kelso et al. 2019). Integrated clones were selected using hygromycin (0.2 ug/uL) and subsequently screened by PCR analysis of the expected 3’ integration site at the FRT locus.

#### Cas9/sgRNA expressing plasmids

The sgRNAs were cloned into the Cas9-expressing plasmid px459 (Addgene 62988, deposited by Dr. Feng Zhang (Ran et al. 2013)) by ligation of annealed oligonucleotides into *Bbs*I digested plasmid. Oligonucleotides were purchases from IDT. Each of the 3’ GFP fragment sequences were designed to contain a sgRNA target sequence that could be used to induce a Cas9 break at the edge of the homology. Hence, each 3’gFP reporter cell line contains a unique sgRNA target sequence: sg150 bp (pCMD35A) for 3’ gFP 150 bp Hom, sg50bp (pCMD34A) for 3’ gFP 50 bp Hom and sg0bp (pCMD32A) for 3’ gFP 0 bp. In addition, all 3’ GFP fragments were designed to contain a restriction cutting site of the rare endonuclease I-*Sce*I 12 bp upstream from the sequences. Accordingly, a sgRNA targeting the I-*Sce*I cutting site was cloned (pCMD31A): sgI-*Sce*I. A sgRNA targeting the AAVS1 locus was cloned and used as a control (pCMD36A): sgCTRL.

- pCMD31A (sgISce-I): GGATAACAGGGTAATCCCGG
- pCMD32A (sg0bp): GGCTGAAGCACTGCACGAAT
- pCMD34A (sg50bp): GGCACGGGCAGCTTGCCAAT
- pCMD35A (sg150bp): TTACGTCGCCGTCCAGCAAT
- pCMD36a (sgCTRL): GAGCCACATTAACCGGCCCT

All sgRNA were validated for targeting Cas9 by assaying the percentage of insertion and deletion mutations at the intended locations using TIDE analysis (Brinkman et al. 2014). For this, we generated amplicons from genomic DNA of each reporter cell line transfected with the respective Cas9/sgRNA plasmid using primer pcDNA5Fw (5’-TACCATGGTGATGCGGTTTT-3’) and primer GFPRev (5’-CGGCCATGATATAGACGTTG-3’) amplified with Platinum HiFi Supermix (ThermoFisher). Untransfected cells were used as a reference control. PCR products were then Sanger sequencing and analyzed using TIDE analysis to quantify the frequency of mutagenesiss.

### L1 retrotransposition GFP reporter assay

#### Transfection-based

The L1 GFP reporter assay for retrotransposition was performed in the parental U2OS cells as previously described (Kopera et al. 2016; Mendez-Dorantes et al. 2024) with the following modifications. Briefly, 0.5 x10^5^ cells were seeded in 12-well plates (d1). The following day cells were transfected using FugeneHD (Promega) with 200 ng of sgCTRL plasmid (pCMD36A) with 400 ng of the pCEP4-Puro L1-GFP reporter plasmids (d 2): pCMD66B (WT), pCMD67A (D702Y, RTmut) and pCMD68A (H230A, ENmut). In addition, equivalent amount of a pCEP4-Puromycin vector expressing a GFP cassette (MT498, pCEP4-GFP) and the Cas9/sgCTRL plasmid was transfected in parallel to monitor transfection efficiency by assaying the percentage of GFP+ cells on d4 by flow cytometry (BD LSRFortessa). For cells transfected with the L1 plasmids, media was changed 12 hours after transfection (d3) and then again the following day (d4) to be supplemented with media containing 1 ug/ mL of Puromycin. Cells were then incubated until d8 when cells were collected and assayed for the percentage of GFP+ cells by flow cytometry (BD LSRFortessa). Singlets were gated on side-scatter versus forward scatter and GFP+ cells were gated on GFP versus autofluorescence (PE). Cells with autofluorescence were detected on a diagonal line, whereas cells showing increased green fluorescence (GFP+) were gated above the autofluorescence diagonal line. We normalized the percentage of GFP+ cells from the cells transfected with the L1 reporter assay to the percentage of GFP+ cells from cells transfected with the GFP plasmid.

#### Tet-On L1-GFP reporter cell line

We also used a Tet-On ORFeusHs-GFP HeLa reporter cell line that we developed to assay frequency of L1 retrotransposition. To establish this reporter cell line, we exploited the Sleeping Beauty DNA transposon system to stably deliver a Tet-On L1 (ORFeusHs) cassette that contains GFP-AI cassette. For this, we transfected HeLa cells using Viafect (Progema) with an expression vector for the Sleepy Beauty transposase (pCMV(CAT)T7-SB100), and the donor plasmid containing Sleepy Beauty inverted terminal repeats flanking the Tet-On ORFeusHs-GFP cassette, and a constitutively expressing rtTA-T2A-Puro-T2A-BFP cassette (pCMD125cx). Cells were selected with Puro, and single cells were sorted to generate a monoclonal cell line (Tet-On ORFeusHs-GFP HeLa cells), which we confirmed via immunoblotting of the L1-encoded proteins and induction GFP+ cells after treatment with doxycycline (1ug/mL). For experiments with siRNA experiments, forward transfection was first performed of 15nM siRNA at a cell density of 1x10^5^ cells per well in 12-well plates using Lipofectamine RNAiMAX. The following day cells were then washed and then fed with media containing doxycycline (1ug/mL) to induce the L1 reporter. GFP+ cells were analyzed as a mentioned above. To deplete BRCA1 or CtIP, we used SMARTpool ON-TARGETplus siRNAs targeting BRCA1 (L-003461-00-0005, siBRCA1) or CtIP (L-011376-00, siCtIP) and ON-TARGETplus non-targeting siRNA control #1 (D-001810-01-20, siCTRL).

### GFP recombination reporter assay for L1 retrotransposition-mediated chromosomal rearrangements

#### GFP Recombination assay between L1 intermediates and Cas9-mediated DNA breaks

Each 3’gFP reporter cell line was plated at a cell density of 0.5 x10^5^ cells per well in 12-well plates (d1). The following day cells were transfected using FugeneHD (Promega) with 200 ng of the indicated Cas9/ sgRNA plasmid and 400 ng of each of the pCEP4-Puro L1-5’ GFp reporter plasmids (d2): pCMD69A (WT), pCMD70C (RTmut, D702Y) or pCMD71B (ENmut, H230A). In addition, equivalent amount of the pCEP4-Puromycin vector expressing a GFP cassette (MT498, pCEP4-GFP) and the Cas9/sgCTRL plasmid was transfected in parallel to monitor transfection efficiency by assaying the percentage of GFP+ cells on d4 by flow cytometry (BD LSRFortessa). For cells transfected with the L1 plasmids, media was changed 12 hours after transfection (d3) and then again the following day (d4) to be supplemented with media containing 1 ug/ mL of Puromycin. Cells were then incubated until d8 when cells were collected and assayed for the percentage of GFP+ cells by flow cytometry (BD LSRFortessa). Singlets were gated on side-scatter versus forward scatter and GFP+ cells were gated on GFP versus autofluorescence (PE). Cells with autofluorescence were detected on a diagonal line, whereas cells showing increased green fluorescence (GFP+) were gated above the autofluorescence diagonal line. The percentage of GFP+ cells from the reporter assays were normalized relative to the percentage of GFP+ cells from cells transfected with the GFP plasmid.

#### GFP Recombination assay between L1 insertion intermediates

Parental U2OS cells were plated at a cell density of 1x10^5^ cells per well in 12-well plates (d1). The following day cells were transfected using FugeneHD (Promega) with 600 ng each of the pCEP4-Puro L1-5’ GFp reporter plasmids, pCMD69A (WT), pCMD70C (RTmut, D702Y) or pCMD71B (ENmut, H230A), plus 600 ng of each of the pCEP4-Puro L1-3’gFP reporter plasmids (L1-3’gFP-150bp Hom or L1-3’gFP-50bp Hom). In addition, a pCEP4-Puromycin vector expressing a GFP cassette (MT498, pCEP4-GFP) and the respective pCEP4-Puro L1-3’gFP reporter plasmid were transfected in parallel to monitor transfection efficiency by assaying the percentage of GFP+ cells. For all cells transfected, media was changed 12 hours after transfection (d3). Cells were then incubated until d6 when cells were collected and assayed for the percentage of GFP+ cells by flow cytometry (BD LSRFortessa). Singlets were gated on side-scatter versus forward scatter and GFP+ cells were gated on GFP versus autofluorescence (PE). Cells with autofluorescence were detected on a diagonal line, whereas cells showing increased green fluorescence (GFP+) were gated above the autofluorescence diagonal line. The percentage of GFP+ cells from the reporter assays were normalized relative to the percentage of GFP+ cells from cells transfected with the GFP plasmid.

#### siRNA experiments

For experiments with siRNA experiments, forward transfection was first performed of 15nM siRNA (siCTRL, siBRCA1, or siCtIP) at a cell density of 1x10^5^ cells per well in 12-well plates using Lipofectamine RNAiMAX. The following day cells were then washed and either of the protocols above were performed.

### HDR-DR-GFP reporter assay

HDR reporter assay DR-GFP U2OS cells were obtained from Dr. Jeremy Stark and cultured in high glucose DMEM supplemented with 10% FBS. The DR-GFP reporter assay was performed as previously described (Gunn and Stark 2012). Briefly, forward transfection was first performed of 15nM siRNA (siCTRL, siBRCA1 or siCtIP) at a cell density of 1x10^5^ cells per well in 12-well plates using Lipofectamine RNAiMAX. The following day cells were then washed and transfected with 600 ng of pCAGGS-I-SceI or pCAGGS-NZE-GFP to monitor transfection efficiency. The following day cells were washed and fed with fresh media. Cells were incubated for four days and analyzed by flow cytometry (BD LSRFortessa). Singlets were gated on side-scatter versus forward scatter and GFP+ cells were gated on GFP versus autofluorescence (PE). Cells with autofluorescence were detected on a diagonal line, whereas cells showing increased green fluorescence (GFP+) were gated above the autofluorescence diagonal line. The percentage of GFP+ cells from the reporter assays were normalized relative to the percentage of GFP+ cells from cells transfected with the GFP plasmid for each siRNA condition.

### Immunoblotting

To immunoblot for L1 encoded ORF1p and ORF2p, 0.3 million cells from the parental U2OS cell line were seeded in two mL of regular media per well of a 6-well plate. The next day, cells were fed with 2 mL of antibiotic-free media and transfected with 1200 ng of the indicated pCEP4-Puro plasmid using FuGENE HD Transfection reagent. The following day, cells were fed with fresh regular media containing puromycin (1 ug/mL) to select for transfected cells. Importantly, one well of cells that were untransfected (UN) we used as a negative control. Cells were collected 1 day post Puromycin selection for protein extraction. Protein was extracted from cells using radioimmunoprecipitation assay buffer (Boston BioProducts BP-115) supplemented with protease and phosphatase inhibitors (Cell Signaling, 5872S). Gel electrophoresis was performed on protein extracts using 4 to 20% Mini-PROTEAN TGX gels (Bio-Rad, 456-1095). Proteins were then transferred to low fluorescence polyvinylidene difluoride membranes using Trans-Blot Turbo (Bio-Rad). Membranes were blocked using EveryBlot Blocking Buffer (Bio-Rad, 12010020) or Intercept Blocking Buffer (Li-COR, 927-60001), and probed with primary ORF2p (abcam, ab263071), ORF1p (Millipore Sigma, MABC1152), beta-tubulin (Cell Signaling, 2128S), followed by secondary antibodies (IRDye 800CW goat anti-mouse IgG, 925-32210; IRDye 680RD goat anti-rabbit IgG, 925-68071; and anti-rabbit IgG horseradish peroxidase (HRP), 7074S). ECL substrate (Thermo Scientific, 34580) was used to develop HRP signals. Immunoblotting signals were detected using the ChemiDoc imaging system (Bio-Rad).

### Targeted Cas9 ONT sequencing

150bpHom-3’ gFP U2OS cells were co-transfected with the L1-5’GFp plasmid and the Cas9/sg150bp plasmid to induce GFP+ cells, as described above. GFP+ were collected seven days after transfection and sorted using BD Aria III SORP (DFCI Flow Cytometry Core). Sorted cells were cultured in DMEM with 20% FBS for the initial 7 days to facilitate cell recovery. GFP+ cells were expanded, and high molecular weight (HMW) genomic DNA was extracted using phenol: chloroform extraction. Genomic DNA was first analyzed for the predicted GFP recombination sequence using PCR analysis with primer hPGKFw (5’-TATTTATGCAGAGGCCGAGG-3’) and primer GFPRev (5’-CGGCCATGATATAGACGTTG-3’) amplified with Platinum HiFi Supermix (ThermoFisher). PCR product was validated using Sanger sequencing. Cas9 targeted nanopore sequencing (McDonald et al. 2021) was performed on the HMW genomic DNA to sequence the GFP recombination events. 5 μg of HMW genomic DNA was fragmented to 30 kb using Megaruptor 3 (Diagenode) and DNA ends were dephosphorylated using recombinant shrimp alkaline phosphatase (rSAP, NEB M0371). Dephosphorylated DNA fragments were incubated with pre-assembled ribonucleoproteins of *S. pyogenes* Cas9 nuclease (NEB M0386) and synthesized sgRNAs (IDT) targeting the 3’ end of GFP: TTCAAGTACCGCCATGCCCGA (sgGFP_A) and GTCGCCCTCGAACTTCACCT (sgGFP_B). Importantly, both sgRNA were designed to target the 3’ GFP sequence inserted at the chromosomal FRT locus to exclude sequencing of canonical L1 insertions with incomplete 5’ GFP sequences and to enrich sequencing of GFP recombination sequences at the FRT locus. Cleaved genomic DNA was then ligated to the nanopore sequencing adapter to make a sequencing library with the Ligation Sequencing Kit SQK-LSK114 (Oxford Nanopore Technologies). The library was sequenced on a MinIon flow cell. The sequencing data were base-called using Dorado super-accurate basecalling v4.3.0, 400bps with modified basecaling 5hmC & 5mC (CG contexts) (Oxford Nanopore Technologies). Nanopore reads were mapped using minimap2 with map-ont presets and a fasta file containing the predicted GFP recombination sequence including SVpolyASignal(-)-hPGKpromoter (+)-GFP(+)-bGHpolyASignal(+) to filter out reads containing GFP sequences. Nanopore reads that that mapped to the 5’GFP side of the reference were considered putative rearrangements since these contained the L1 inserted sequence by the L1-5’GFp reporter plasmid, and the rearrangement junction between the 5’GFP sequence and the 3’GFP sequence. In contrast, nanopore reads that mapped to the 3’ GFP side of the reference were considered reads mapping to the chromosomal location where the FRT construct was introduced into U2OS cells. To identify the genomic locations of long reads, these filtered GFP-containing reads were then mapped using minimap2 and a fasta file containing the T2T human reference genome (T2T-CHM13/hs1) and the predicted GFP recombination sequence as an additional chromosome. Additionally, manual curation of the identified reads for known L1-mediated retrotransposition features such as target-site duplications, L1 endonuclease cutting motif sequences, and inserted polyA sequences at the 3’ end of insertions.

### ONT WGS

Parental U2OS cells were transfected with the L1-5’GFp plasmid and the L1-3’gFP plasmid to induce GFP+ cells, as described above. GFP+ cells were collected six days after transfection and sorted via BD Aria III SORP (DFCI Flow Cytometry Core). GFP+ cells were expanded, and genomic DNA was extracted using the Invitrogen PureLink Genomic DNA Mini kit (K182001). Genomic DNA was first analyzed for the predicted GFP recombination sequence using PCR analysis with primer hPGKFw (5-TATTTATGCAGAGGCCGAGG-3’) and primer bGHRev (5’-TAGAAGGCACAGTCGAGG-3’) amplified with Platinum HiFi Supermix (ThermoFisher). PCR product was validated using Sanger Sequencing. ONT WGS was then performed on the genomic DNA to determine the genomic locations of the GFP recombination events (Jain et al. 2018). A total of 6 μg of extracted genomic DNA was fragmented to 30 kb using the Megaruptor 3 (Diagenode). Fragmented genomic DNA was then ligated to the nanopore sequencing adapter to make a sequencing library with the Ligation Sequencing Kit SQK-LSK114 (Oxford Nanopore Technologies). 300 ng of library was sequenced on the PromethION flowcell FLO-PRO114M using the PromethION Solo device for 3 d with 2 washes and reloads of 300ng of library after 24 and 48 hours using the Flow Cell Wash Kit EXP-WSH004. The sequencing data were base-called using Dorado super-accurate basecalling v4.3.0, 400bps with modified basecaling 5hmC & 5mC (CG contexts) (Oxford Nanopore Technologies). Nanopore reads were mapped using minimap2 with map-ont presets and a fasta file containing the predicted GFP recombination sequence including SVpolyASignal(-)-hPGKpromoter (+)-GFP(+)-bGHpolyASignal(+) to filter out reads containing GFP sequences. Nanopore reads that spanned the complete predicted GFP recombination sequence were designated as putative rearrangements, whereas nanopore reads that mapped to the 5’GFP side of the reference were considered putative canonical insertions generated by the L1-5’GFp reporter plasmid and the nanopore reads that mapped to the 3’GFP side of the reference were considered putative canonical insertions generated by the L1-3’gFP reporter plasmid. To identify the genomic locations of these L1 inserted sequences, filtered GFP-containing reads were then mapped using minimap2 and a fasta file containing the T2T human reference genome (T2T-CHM13/hs1) and the predicted GFP recombination sequence as an additional chromosome. Additionally, manual curation of the identified reads for known L1-mediated retrotransposition features such as target-site duplications, L1 endonuclease cutting motif sequences, and inserted polyA sequences at the 3’ end of insertions.

## Data Availability

DNA sequencing data will be deposited to the Sequencing Read Archive (SRA).

## Author Contributions

C.M.-D. and K.H.B. conceived the project and designed the experiments. C.M.-D., J.C.K. and P.S. performed all the experiments and data analyses. C.-T.L. performed Cas9 targeted nanopore sequencing and J.C.K. and A.B. performed nanopore whole-genome sequencing. C.-T.L. and A.B. filtered all the GFP-containing long reads and C.M.D. analyzed the L1-mediated rearrangements and insertions. C.M.-D. wrote the manuscript and C.M.-D. and K.H.B. revised the manuscript with input from all authors.

## Acknowledgements

K.H.B. is supported by the National Institutes of Health (R01CA240816, R01CA276112, R01CA289390, UG3NS132127) and by the Dana-Farber Cancer Institute (Innovations Research Fund). C.-T.L. is supported by an American Cancer Society Postdoctoral Fellowship Award. C.M.-D. is a fellow of The Jane Coffin Childs Fund for Medical Research, a recipient of the Charles A. King Trust Postdoctoral Research Fellowship Award, and a Forbeck Scholar. Some diagrams were created using BioRender.com.

---

**Figure S1.**
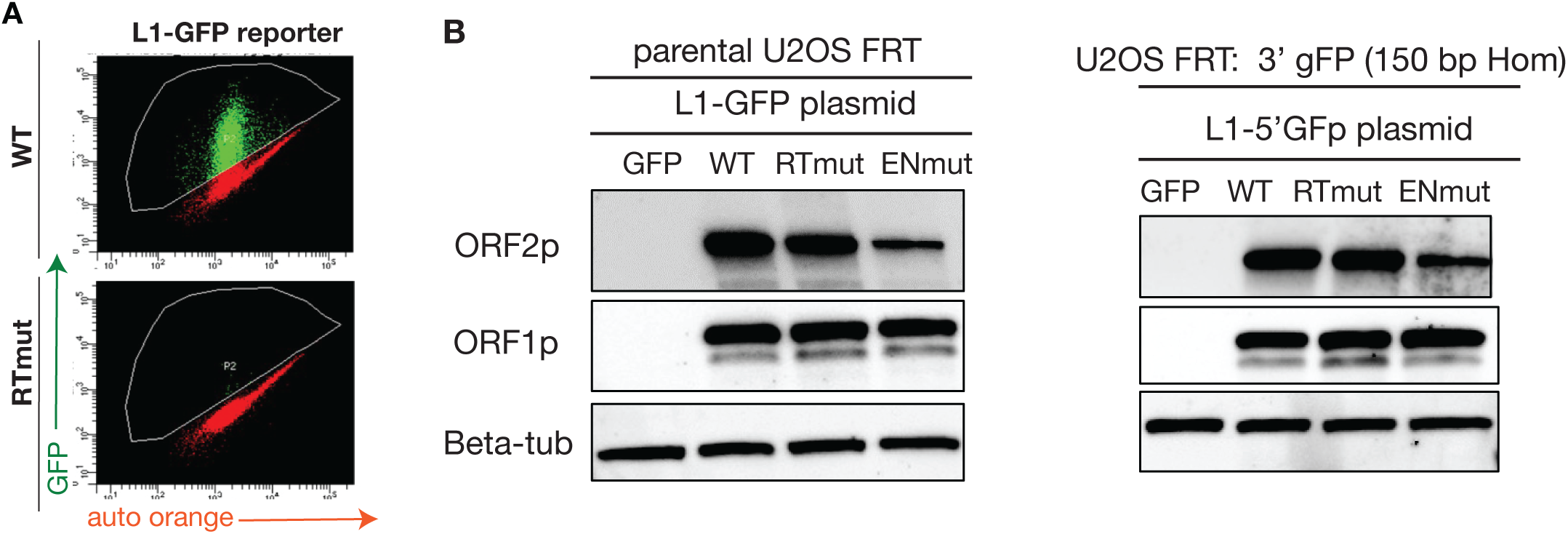
Validation of the L1-5’GFp reporter plasmid. **A** Shown are representative flow cytometry plots of U2OS cells transfected with WT or reverse transcriptase mutant (RTmut, D702Y) L1-GFP reporter. **B.** Shown are representative immunoblots showing similar ORF2p and ORF1p expression from the classic L1-GFP plasmid and the new modified L1-5’ GFp plasmid.

**Figure S2.**
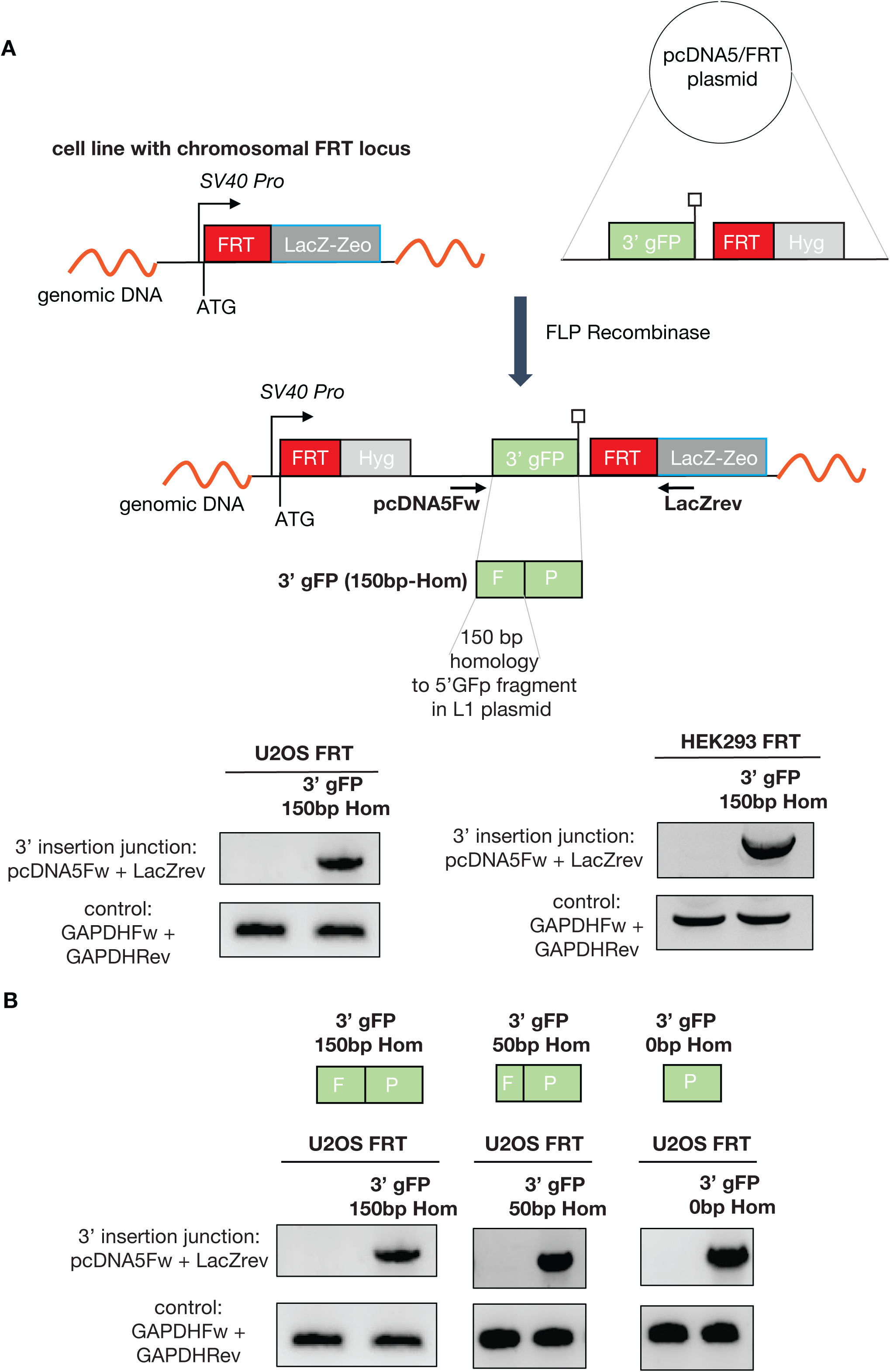
Validation of cell engineering of U2OS and HEK293 cells containing 3’gFP fragments at a chromosomal FRT locus. **A**. Shown is a schematic of the FRT/Flp system used to introduce 3’gFP fragments into a specific chromosomal FRT locus in U2OS or HEK293 cells. Briefly, a 3’gFP fragment containing 150 bp homology (3’gFP 150bp Hom) to the 5’GFp fragment included in the L1-5’GFp plasmid was synthesized and cloned into the pcDNA5/FRT plasmid. The 3’gFP 150bp Hom construct was introduced into U2OS or HEK293 cells with a chromosomal FRT locus using FLP recombinase. Targeted clones are selected under hygromycin and screened for the integrated 3’gFP fragment using PCR primers flanking the 3’ FRT integration site: pcDNA5Fw (L1 5’GFp plasmid primer) and LacZrev. Shown are PCR amplification products from U2OS FRT 3’ gFP (150bp Hom) cells or HEK293 FRT 3’ gFP (150bp Hom) using primers flanking the 3’ FRT integration site: pcDNA5Fw and LacZrev. The respective parental cells were used as a negative control. Amplification of GAPDH locus was used as a positive control. **B.** Two additional U2OS reporter cell lines were established containing 3’gFP fragments with varying degrees of homology to the 5’ GFp fragment from the L1-5’GFp plasmid. Namely, 3’ gFP fragments containing either 50 bp homology (3’gFP 50bp Hom) or 0 bp homology (3’ gFP 0bp Hom) were cloned into the pcDNA5/FRT plasmid and introduced into U2OS cells with a chromosomal FRT locus using the FRT/Flp recombinase system. Shown are PCR amplification products from U2OS FRT 3’ gFP (50bp Hom) cells or U2OS FRT 3’ gFP (50bp Hom) cells using primers flanking the 3’ FRT integration site: pcDNA5Fw and LacZrev. The respective parental cells were used as a negative control. Amplification of GAPDH locus was used as a positive control.

**Figure S3.**
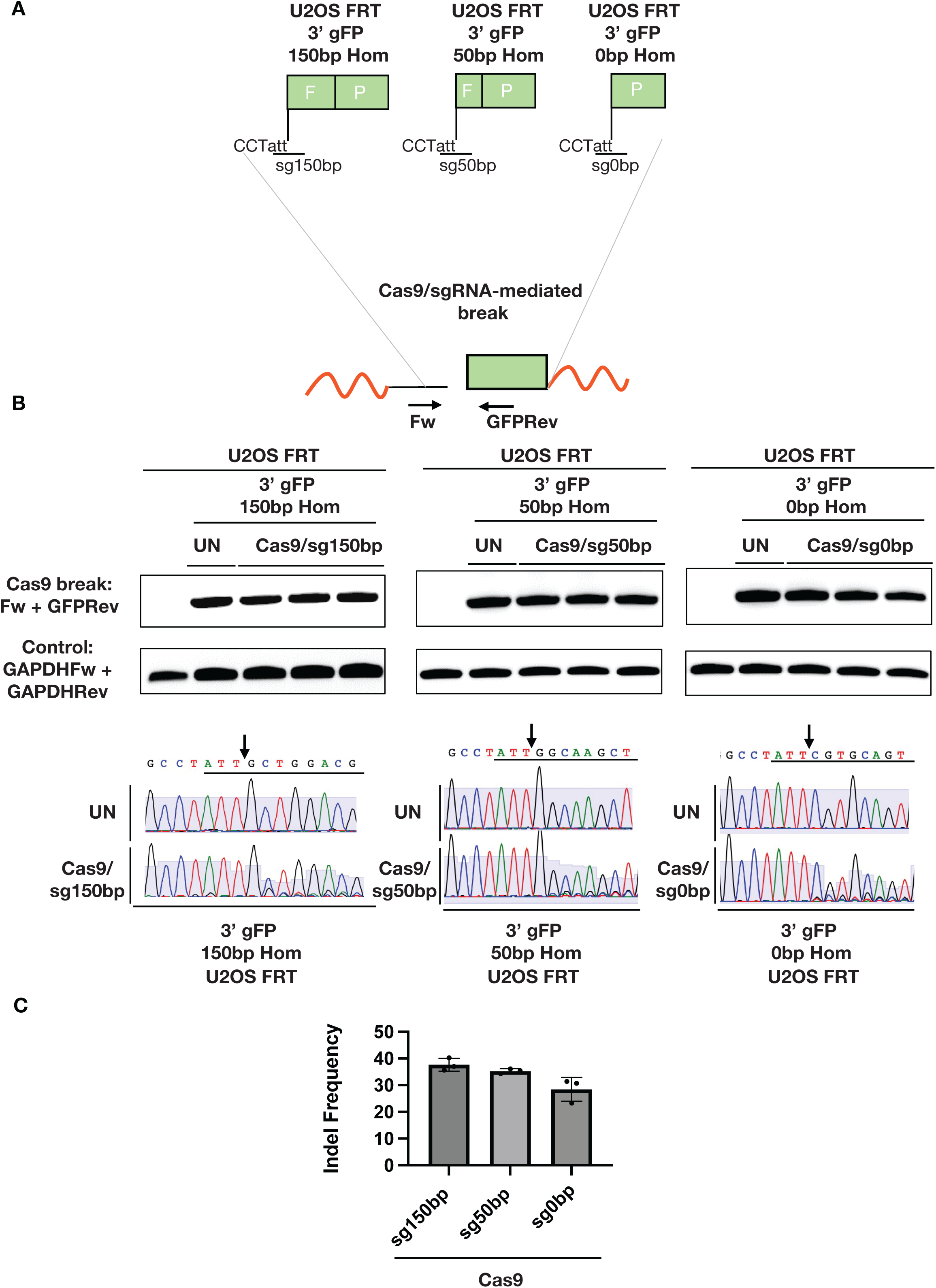
Validation of Cas9 cutting at the predicted genomic locus at the edge of the integrated 3’ GFP sequences. **A.** Each 3’ gFP integrated at the chromosomal FRT locus was designed to contain a sgRNA target sequence to induce a Cas9-mediated DNA double-strand break at the edge of the 3’GFP sequences as depicted in the diagram. An amplicon can be generated flanking the targeted Cas9/sgRNA-mediated DNA double-strand break to assay the frequency of insertion and deletion mutations (indel). **B.** Shown are representative amplicons generated from each 3’ gFP reporter cell line transfected with the respective Cas9/sgRNA expressing plasmid. Amplicons were generated using primers flanking the Cas9 breaks: pDNA5Fw and GFPRev. The respective parental cells were used as a negative control. Amplification of the GAPDH locus was used as a positive control. Also, shown are the Sanger sequencing traces from the amplicons showing mutagenesis at the Cas9 targeted sites in the transfected cells but not in the untransfected cells. **C.** Shown are the indel frequency at the targeted sites in each 3’ gFP reporter cell line after transfection with the respective Cas9/sgRNA expressing plasmids. Indel frequency was assayed using TIDE (Tracking of Indel By Decomposition) analysis of the amplicons generated after the induction of the targeted Cas9 breaks. Amplicons derived from untransfected cells were used as a reference for the TIDE analysis.

**Figure S4.**
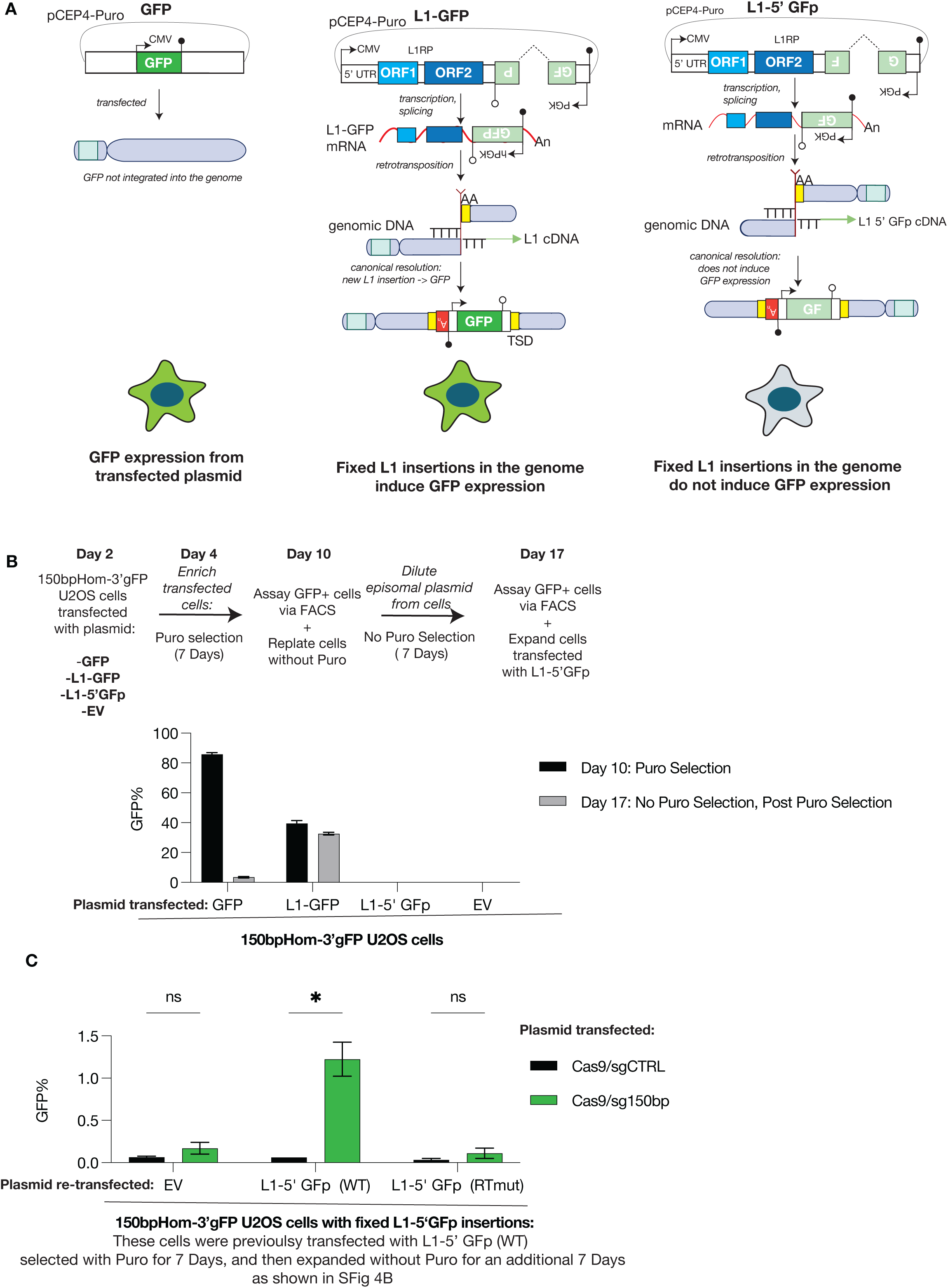
Sequential expression of the L1-5’GFp plasmid and the Cas9/150bp expressing plasmid fails to induce GFP+ cells. **A.** A schematic showing the GFP status of cells transfected with a pCEP4-Puro plasmid expressing GFP, L1-GFP or L1-5’GFp. *Left. GFP:* GFP expression is induced from the GFP transgene expressed from the plasmid transfected in cells without integrating into the genome. *Middle. L1-GFP:* GFP expression is induced from fixed L1 insertions in the genome generated via L1 retrotranspostion in cells transfected with the classic L1-GFP retrotransposition reporter plasmid. *Right. L1-5’GFp:* GFP expression is not induced with fixed L1 insertions that contain an incomplete 5’GFP fragment generated via L1 retrotranspositon in the genome of cells transfected from the L1-5’GFp reporter plasmid. **B.** Shown is the percentage of GFP+ cells of 150bpHom-3’gFP U2OS cells transfected with a pCEP4-Puro plasmid expressing GFP, L1-GFP or L1-5’GFp and selected with Puro for 7 days, as well after continued culture without Puro for an additional 7 days. Cells transfected with GFP alone are about 90% GFP+ after 7 days of Puro selection but become nearly 0% GFP+ after continued culture without Puro showing that the pCEP4 plasmid is diluted from cells. In contrast, the percentage of GFP+ cells remains similar for cells transfected with the classic L1-GFP reporter plasmid since GFP expression results from L1 insertions fixed in the genome. The prediction is that cells transfected with the L1-5’GFp reporter plasmid will also contain fixed L1 insertions containing 5’ GFP fragments. **C.** Shown is the frequency of GFP+ cells from 150bpHom-3’gFP U2OS cells that were previously transfected with the WT L1-5’GFp plasmid as described in SFig 4B and then <u>again</u> co-transfected with an EV, or WT or RTmut L1-5’GFp plasmid and the Cas9/sgCTRL or the Cas9/sg150bp plasmid. n=3. P-value calculated using unpaired two-tailed Student’s t tests with the Holm-Sidak correction for multiple comparisons.

**Figure S5.**
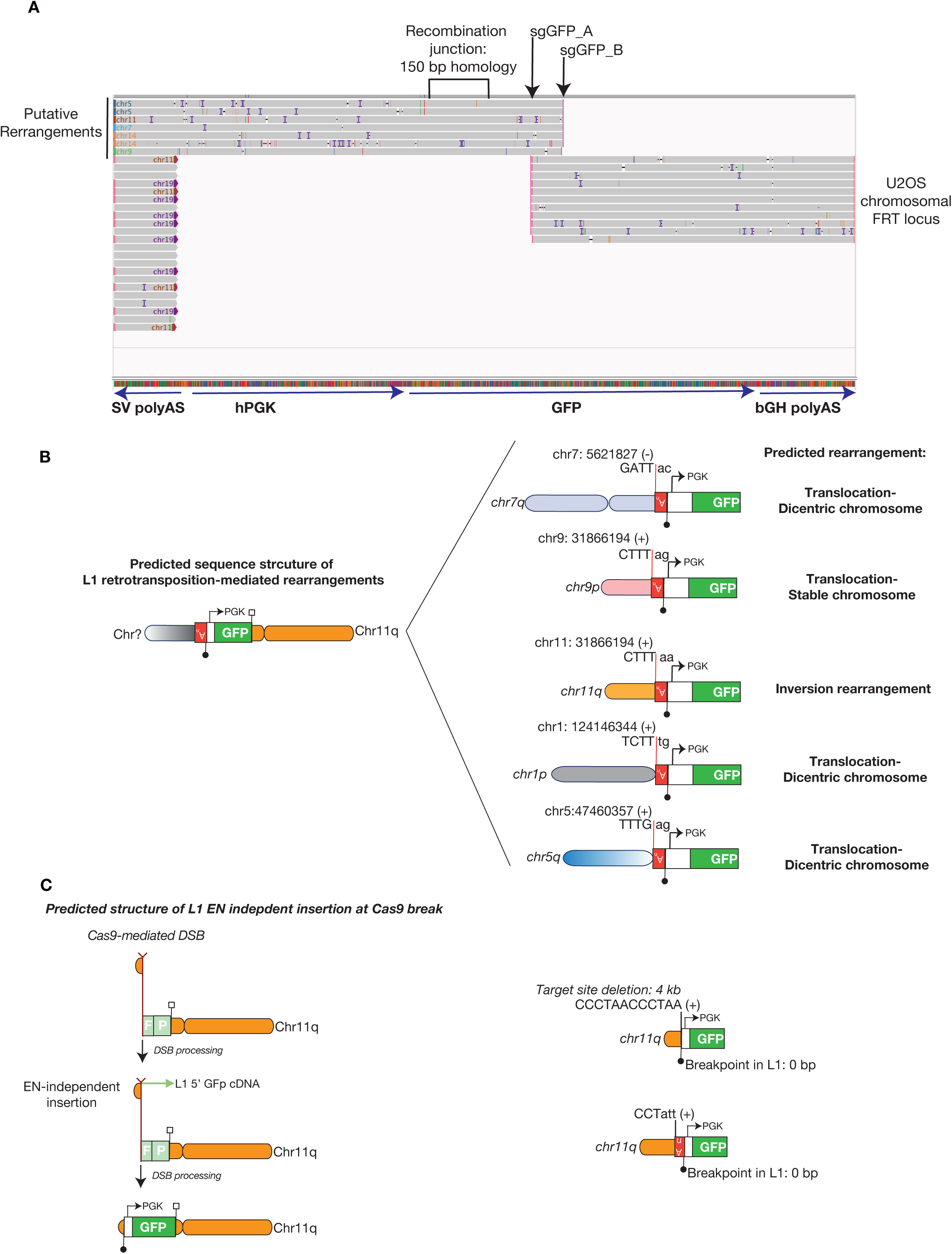
Cas9 targeted ONT sequencing captures L1-mediated rearrangements, and L1 endonuclease-independent insertions at the induced chromosomal break. **A.** Shown are representative long reads derived from Cas9 targeted ONT sequencing of GFP+ cells mapped to the predicted GFP recombination reference sequence. Reads mapping to the right of the reference sequence map to the 3’ end of GFP and to chromosome 11 (7784 +) where the FRT construct was integrated. Reads multi-map to the 5’ side of the reference because the FRT construct also contains a SVpolyA sequence. **B.** Shown are schematics of the predicted sequence structure of the L1 retrotransposition-mediated rearrangements showing evidence of L1 TPRT activity at the genomic junction. **C.** Shown are schematics of the sequence structure of the L1 EN-independent insertions at the Cas9-mediated chromosomal break at the FRT locus in chromosome 11q.

**Figure S6.**
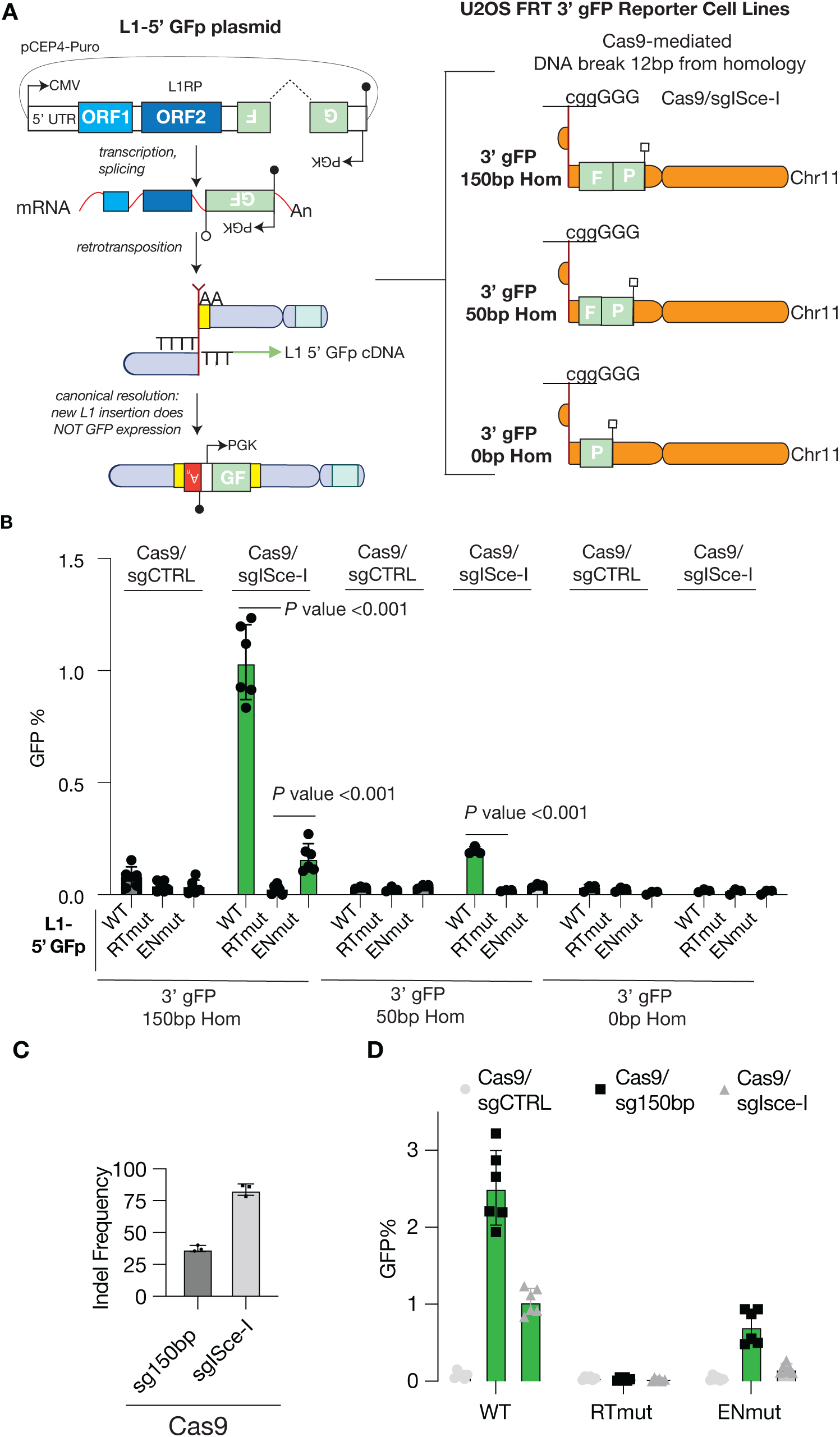
Influence of the proximity of the chromosomal break next to the homology of 3’GFP sequences on L1 retrotransposition-mediated rearrangements. **A.** Shown are schematic of the GFP reporter assays modeling the recombination between L1 retrotransposition cDNA products and chromosomal break ends containing varying degree of shared homology similar as in Figure 3A. However, the inducing chromosomal break in all reporter cell lines is generated using the same Cas9/sgRNA plasmid targeting an I-SceI cutting site, which located 12 bp away from the 3’ GFP sequences in all reporter cell lines. **B.** Shown is the GFP frequency induced in each U2OS 3’gFP reporter cell line transfected with WT, RTmut or ENmut L1-5’ GFp reporter plasmid, and the Cas9/sgSce-I expressing plasmid or the control Cas9/sgCTRL expressing plasmid. n = 6-3. P-value calculated by comparing WT and ENmut versus RTmut in each condition using a two-way ANOVA with the Dunnett correction for multiple comparisons. **C.** Shown are the indel frequencies at the targeted sites in the 150bpHom-3’ gFP U2OS cell line after transfection with the Cas9/sg150bp expressing plasmid or the Cas9/sgISce-I expressing plasmid. Indel frequency was assayed using TIDE (Tracking of Indel By Decomposition) analysis of the amplicons generated after the induction of the targeted Cas9 breaks. Amplicons derived untransfected cells were used as a reference control for TIDE analysis. **D.** GFP frequencies induced in 150bp-Hom-3’ gFP U2OS cells transfected with WT, RTmut or ENmut L1-5’ GFp reporter plasmid, and Cas9/sgCTRL, Cas9/sg150bp, or Cas9/sgISce-I expressing plasmid. n = 6.

**Figure S7.**
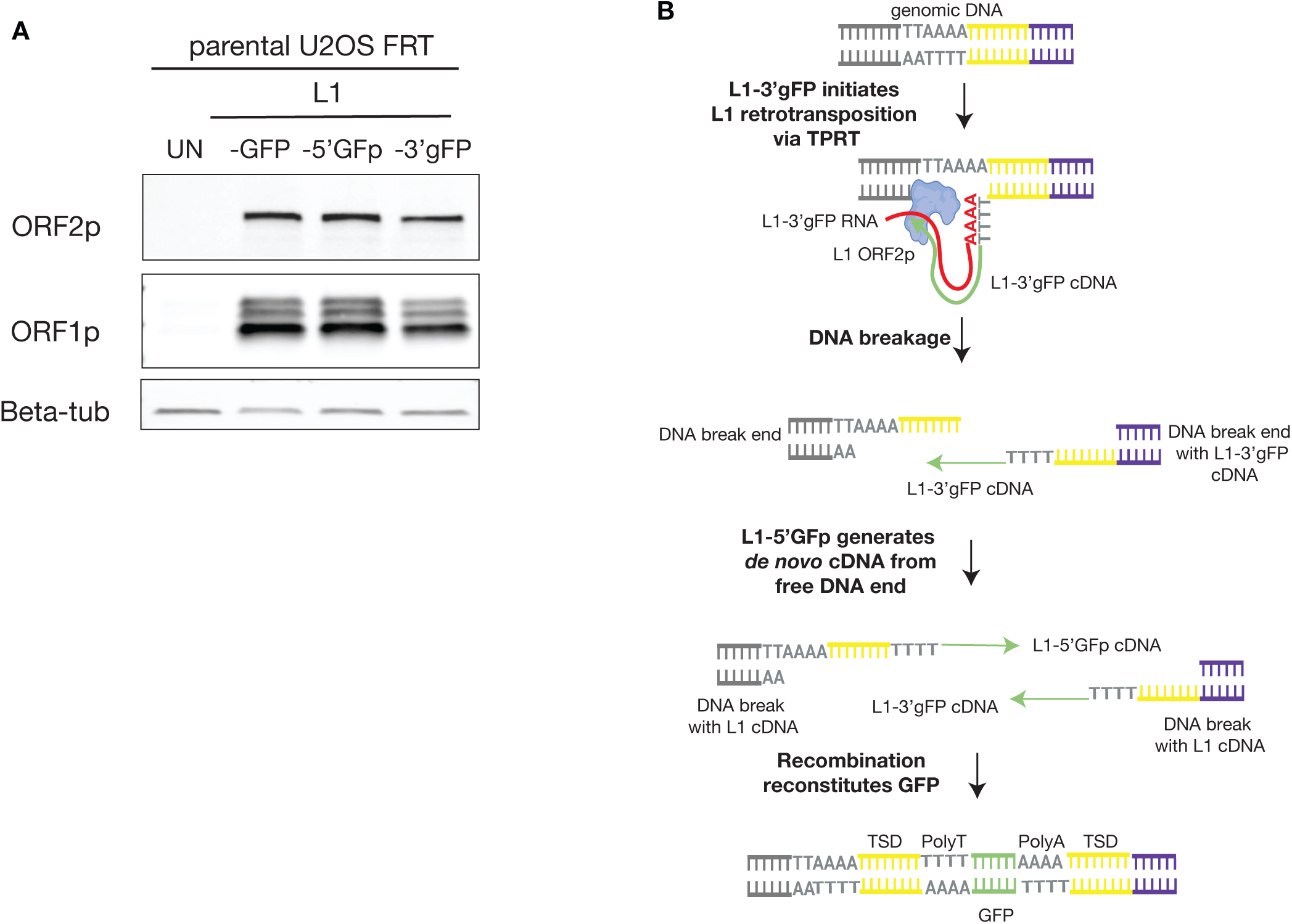
Validation of L1-3’ gFP reporter plasmid. **A)** Representative immunoblots showing similar ORF1p and ORF2p expression levels from the classic L1-GFP reporter plasmid, and the new L1-3’ gFP reporter plasmid. **B)** A schematic showing a working model how two RT reactions on the same chromosomal location can reconstitute GFP via recombination (n=2):) L1-3’gFP initiates TPRT to generate the 3’GFP cDNA product and results in DNA breakage and 2) L1-5’GFp generates 5’GFP cDNA product from the free DNA break and the two cDNA products recombine to seal the insertion in the genome (SFig 7B, Table 2). This model is supported by the fact that both two events contained target site duplications and inserted polyA sequences

**Figure S8.**
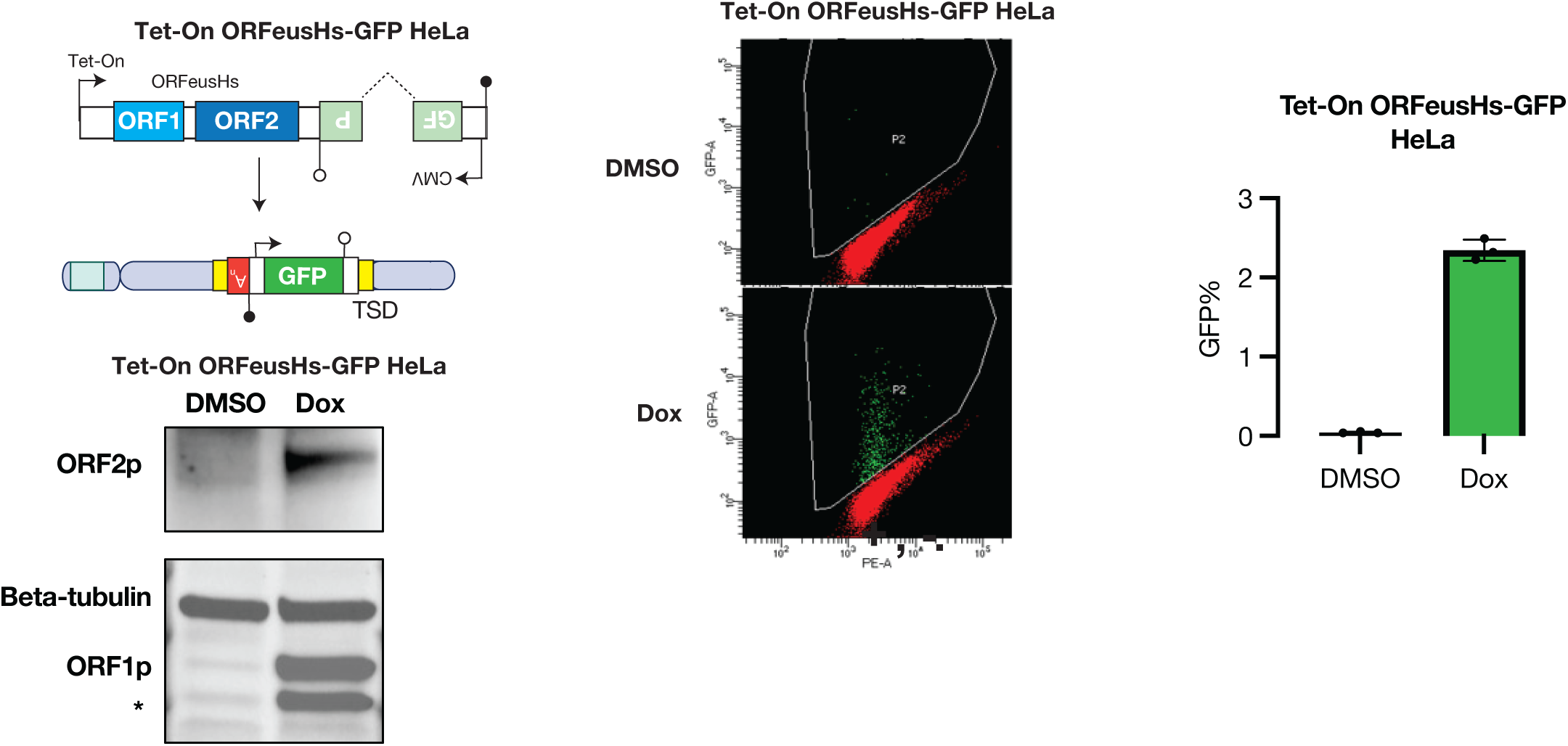
Validation of Tet-On ORFeusHs-GFP Hela reporter cell line. Shown is a schematic of the Tet-On ORFeusHs-GFP reporter construct that upon doxycycline treatment results in de novo L1 insertions inducing GFP expression. Also, shown are immunoblotting analysis showing that only after doxycycline treatment L1 encoded proteins are expressed in HeLa cells. Moreover, representative flow cytometry plots and the frequency of of GFP+ cells of the Tet-On ORFeusHs-GFP HeLa reporter cell line after treatment with DMSO as a control versus doxycycline.

## Reference

Ardeljan D, Steranka JP, Liu C, Li Z, Taylor MS, Payer LM, Gorbounov M, Sarnecki JS, Deshpande V, Hruban RH et al. 2020. Cell fitness screens reveal a conflict between LINE-1 retrotransposition and DNA replication. Nature structural & molecular biology 27: 168–178.

Baldwin ET, van Eeuwen T, Hoyos D, Zalevsky A, Tchesnokov EP, Sánchez R, Miller BD, Di Stefano LH, Ruiz FX, Hancock M et al. 2024. Structures, functions and adaptations of the human LINE-1 ORF2 protein. Nature 626: 194–206.

Bhargava R, Onyango DO, Stark JM. 2016. Regulation of Single-Strand Annealing and its Role in Genome Maintenance. Trends in genetics : TIG 32: 566–575.

Brinkman EK, Chen T, Amendola M, van Steensel B. 2014. Easy quantitative assessment of genome editing by sequence trace decomposition. Nucleic acids research 42: e168.

Cajuso T, Sulo P, Tanskanen T, Katainen R, Taira A, Hänninen UA, Kondelin J, Forsström L, Välimäki N, Aavikko M et al. 2019. Retrotransposon insertions can initiate colorectal cancer and are associated with poor survival. Nature communications 10: 4022.

Feng Q, Moran JV, Kazazian HH, Jr., Boeke JD. 1996. Human L1 retrotransposon encodes a conserved endonuclease required for retrotransposition. Cell 87: 905–916.

Flasch DA, Macia Á, Sánchez L, Ljungman M, Heras SR, García-Pérez JL, Wilson TE, Moran JV. 2019. Genome-wide de novo L1 Retrotransposition Connects Endonuclease Activity with Replication. Cell 177: 837–851.e828.

Gasior SL, Wakeman TP, Xu B, Deininger PL. 2006. The human LINE-1 retrotransposon creates DNA double-strand breaks. Journal of molecular biology 357: 1383–1393.

Gilbert N, Lutz S, Morrish TA, Moran JV. 2005. Multiple fates of L1 retrotransposition intermediates in cultured human cells. Molecular and cellular biology 25: 7780–7795.

Gilbert N, Lutz-Prigge S, Moran JV. 2002. Genomic deletions created upon LINE-1 retrotransposition. Cell 110: 315–325.

Gunn A, Stark JM. 2012. I-SceI-based assays to examine distinct repair outcomes of mammalian chromosomal double strand breaks. *Methods in molecular biology (Clifton*, NJ) 920: 379–391.

Helman E, Lawrence MS, Stewart C, Sougnez C, Getz G, Meyerson M. 2014. Somatic retrotransposition in human cancer revealed by whole-genome and exome sequencing. Genome research 24: 1053–1063.

Hoyt SJ, Storer JM, Hartley GA, Grady PGS, Gershman A, de Lima LG, Limouse C, Halabian R, Wojenski L, Rodriguez M et al. 2022. From telomere to telomere: The transcriptional and epigenetic state of human repeat elements. *Science (New York*, NY) 376: eabk3112.

Jain M, Koren S, Miga KH, Quick J, Rand AC, Sasani TA, Tyson JR, Beggs AD, Dilthey AT, Fiddes IT et al. 2018. Nanopore sequencing and assembly of a human genome with ultra-long reads. Nature biotechnology 36: 338–345.

Kelso AA, Lopezcolorado FW, Bhargava R, Stark JM. 2019. Distinct roles of RAD52 and POLQ in chromosomal break repair and replication stress response. PLoS genetics 15: e1008319.

Khazina E, Truffault V, Buttner R, Schmidt S, Coles M, Weichenrieder O. 2011. Trimeric structure and flexibility of the L1ORF1 protein in human L1 retrotransposition. Nature structural & molecular biology 18: 1006–1014.

Kimberland ML, Divoky V, Prchal J, Schwahn U, Berger W, Kazazian HH, Jr. 1999. Full-length human L1 insertions retain the capacity for high frequency retrotransposition in cultured cells. Human molecular genetics 8: 1557–1560.

Kopera HC, Larson PA, Moldovan JB, Richardson SR, Liu Y, Moran JV. 2016. LINE-1 Cultured Cell Retrotransposition Assay. *Methods in molecular biology (Clifton*, NJ) 1400: 139–156.

Kramara J, Osia B, Malkova A. 2018. Break-Induced Replication: The Where, The Why, and The How. Trends in genetics : TIG 34: 518–531.

Kulpa DA, Moran JV. 2006. Cis-preferential LINE-1 reverse transcriptase activity in ribonucleoprotein particles. Nature structural & molecular biology 13: 655–660.

Lee E, Iskow R, Yang L, Gokcumen O, Haseley P, Luquette LJ, 3rd, Lohr JG, Harris CC, Ding L, Wilson RK et al. 2012. Landscape of somatic retrotransposition in human cancers. Science (New York, NY) 337: 967–971.

Li Y, Roberts ND, Wala JA, Shapira O, Schumacher SE, Kumar K, Khurana E, Waszak S, Korbel JO, Haber JE et al. 2020. Patterns of somatic structural variation in human cancer genomes. Nature 578: 112–121.

Liu N, Lee CH, Swigut T, Grow E, Gu B, Bassik MC, Wysocka J. 2018. Selective silencing of euchromatic L1s revealed by genome-wide screens for L1 regulators. Nature 553: 228–232.

Manoj F, Tai LW, Wang KSM, Kuhlman TE. 2021. Targeted insertion of large genetic payloads using cas directed LINE-1 reverse transcriptase. Sci Rep 11: 23625.

Mathias SL, Scott AF, Kazazian HH, Jr., Boeke JD, Gabriel A. 1991. Reverse transcriptase encoded by a human transposable element. *Science (New York*, NY) 254: 1808–1810.

McDonald TL, Zhou W, Castro CP, Mumm C, Switzenberg JA, Mills RE, Boyle AP. 2021. Cas9 targeted enrichment of mobile elements using nanopore sequencing. Nature communications 12: 3586.

Mendez-Dorantes C, Bhargava R, Stark JM. 2018. Repeat-mediated deletions can be induced by a chromosomal break far from a repeat, but multiple pathways suppress such rearrangements. Genes & development 32: 524–536.

Mendez-Dorantes C, Burns KH. 2023. LINE-1 retrotransposition and its deregulation in cancers: implications for therapeutic opportunities. Genes & development 37: 948–967.

Mendez-Dorantes C, Tsai LJ, Jahanshir E, Lopezcolorado FW, Stark JM. 2020. BLM has Contrary Effects on Repeat-Mediated Deletions, based on the Distance of DNA DSBs to a Repeat and Repeat Divergence. Cell reports 30: 1342–1357.e1344.

Mendez-Dorantes C, Zeng X, Karlow JA, Schofield P, Turner S, Kalinowski J, Denisko D, Lee EA, Burns KH, Zhang C-Z. 2024. Chromosomal rearrangements and instability caused by the LINE-1 retrotransposon. bioRxiv: 2024.2012.2014.628481.

Mita P, Sun X, Fenyö D, Kahler DJ, Li D, Agmon N, Wudzinska A, Keegan S, Bader JS, Yun C et al. 2020. BRCA1 and S phase DNA repair pathways restrict LINE-1 retrotransposition in human cells. Nature structural & molecular biology 27: 179–191.

Morales ME, White TB, Streva VA, DeFreece CB, Hedges DJ, Deininger PL. 2015. The contribution of alu elements to mutagenic DNA double-strand break repair. PLoS genetics 11: e1005016.

Moran JV, Holmes SE, Naas TP, DeBerardinis RJ, Boeke JD, Kazazian HH, Jr. 1996. High frequency retrotransposition in cultured mammalian cells. Cell 87: 917–927.

Morrish TA, Garcia-Perez JL, Stamato TD, Taccioli GE, Sekiguchi J, Moran JV. 2007. Endonuclease-independent LINE-1 retrotransposition at mammalian telomeres. Nature 446: 208–212.

Moynahan ME, Chiu JW, Koller BH, Jasin M. 1999. Brca1 controls homology-directed DNA repair. Molecular cell 4: 511–518.

Ostertag EM, Prak ET, DeBerardinis RJ, Moran JV, Kazazian HH, Jr. 2000. Determination of L1 retrotransposition kinetics in cultured cells. Nucleic acids research 28: 1418–1423.

Ran FA, Hsu PD, Wright J, Agarwala V, Scott DA, Zhang F. 2013. Genome engineering using the CRISPR-Cas9 system. Nat Protoc 8: 2281–2308.

Rodic N, Sharma R, Sharma R, Zampella J, Dai L, Taylor MS, Hruban RH, Iacobuzio-Donahue CA, Maitra A, Torbenson MS et al. 2014. Long interspersed element-1 protein expression is a hallmark of many human cancers. The American journal of pathology 184: 1280–1286.

Rodriguez-Martin B, Alvarez EG, Baez-Ortega A, Zamora J, Supek F, Demeulemeester J, Santamarina M, Ju YS, Temes J, Garcia-Souto D et al. 2020. Pan-cancer analysis of whole genomes identifies driver rearrangements promoted by LINE-1 retrotransposition. Nature genetics 52: 306–319.

Scott EC, Gardner EJ, Masood A, Chuang NT, Vertino PM, Devine SE. 2016. A hot L1 retrotransposon evades somatic repression and initiates human colorectal cancer. Genome research 26: 745–755.

Stark JM, Pierce AJ, Oh J, Pastink A, Jasin M. 2004. Genetic steps of mammalian homologous repair with distinct mutagenic consequences. Molecular and cellular biology 24: 9305–9316.

Symer DE, Connelly C, Szak ST, Caputo EM, Cost GJ, Parmigiani G, Boeke JD. 2002. Human l1 retrotransposition is associated with genetic instability in vivo. Cell 110: 327–338.

Tao J, Wang Q, Mendez-Dorantes C, Burns KH, Chiarle R. 2022. Frequency and mechanisms of LINE-1 retrotransposon insertions at CRISPR/Cas9 sites. Nature communications 13: 3685.

Taylor MS, Wu C, Fridy PC, Zhang SJ, Senussi Y, Wolters JC, Cajuso T, Cheng WC, Heaps JD, Miller BD et al. 2023. Ultrasensitive Detection of Circulating LINE-1 ORF1p as a Specific Multicancer Biomarker. Cancer discovery 13: 2532–2547.

Thawani A, Ariza AJF, Nogales E, Collins K. 2024. Template and target-site recognition by human LINE-1 in retrotransposition. Nature 626: 186–193.

Tubio JMC, Li Y, Ju YS, Martincorena I, Cooke SL, Tojo M, Gundem G, Pipinikas CP, Zamora J, Raine K et al. 2014. Mobile DNA in cancer. Extensive transduction of nonrepetitive DNA mediated by L1 retrotransposition in cancer genomes. Science (New York, NY) 345: 1251343.

Wang Y, Lin RZ, Harris M, Lavayen B, Diwanji N, McCreedy B, Hofmeister R, Getts D. 2025. CRISPR-Enabled Autonomous Transposable Element (CREATE) for RNA-based gene editing and delivery. EMBO Rep 26: 1062–1083.

Wei W, Gilbert N, Ooi SL, Lawler JF, Ostertag EM, Kazazian HH, Boeke JD, Moran JV. 2001. Human L1 retrotransposition: cis preference versus trans complementation. Molecular and cellular biology 21: 1429–1439.

Wilkinson ME, Frangieh CJ, Macrae RK, Zhang F. 2023. Structure of the R2 non-LTR retrotransposon initiating target-primed reverse transcription. *Science (New York*, NY) 380: 301–308.

Zumalave S, Santamarina M, Espasandín NP, Garcia-Souto D, Temes J, Baker TM, Pequeño-Valtierra A, Otero I, Rodríguez-Castro J, Oitabén A et al. 2024. Synchronous L1 retrotransposition events promote chromosomal crossover early in human tumorigenesis. bioRxiv: 2024.2008.2027.596794.

